# Dissociable misconfigurations of striatal functional connectivity profiles in smokers

**DOI:** 10.1101/2022.02.15.480576

**Authors:** Cole Korponay, Elliot A Stein, Thomas J Ross

## Abstract

Corticostriatal circuits are central to reward processing and reinforcement learning functions that often become dysregulated in substance use disorder (SUD) and drive compulsive drug use. Human neuroimaging research seeking to identify how corticostriatal circuits become altered in SUD has primarily focused on evaluating connectivity between cortex-striatum node-pairs. Yet, striatal nodes receive appreciable input from many cortical nodes, and the morphological and electrophysiological properties of striatal nodes dictate that combinational features of their multivariate “connectivity profiles” shape their activity more so than any individual cortical node. Here, we introduce an approach for quantifying and statistically evaluating different types of multivariate connectivity profile configuration differences (i.e., aggregate divergence, rank order arrangement, and entropy shift) that may reflect different forms of circuit plasticity, and apply it to nicotine dependent smokers (n=46) as an exemplar SUD. Foremost, we find evidence of significant connectivity profile misconfiguration throughout much of the striatum, suggesting that prior findings of abnormal connections between individual striatal-cortical node-pairs may only represent the “tip of the iceberg” of corticostriatal circuit alteration in nicotine dependence. Moreover, we find that dorsolateral and ventromedial striatum display distinct types of connectivity profile misconfiguration. Whereas dorsolateral striatum almost exclusively displays abnormal rank order arrangement that is present in both the nicotine sated and acutely abstinent states – indicative of a “trait” misconfiguration – ventromedial striatum almost exclusively displays abnormal aggregate divergence that only manifests during acute abstinence – indicative of a “state” misconfiguration. Further, we identify a unique striatal site in the right caudal ventral putamen that displays multiple forms of connectivity profile misconfiguration, where connections with cognitive processing cortical areas overtake those with motor/premotor control cortical areas as the strongest in the connectivity profile, and acute abstinence significantly strengthens this abnormal arrangement. Moreover, the interactive magnitude of these misconfigurations during acute abstinence is significantly linked to dependence severity. Collectively, the present findings underscore the need for increased examination of connectivity profile misconfigurations as a mechanism of SUD etiology and as a potential guide for identifying therapeutic intervention targets.

## Introduction

Nicotine dependence remains the largest cause of preventable death in the United States[1], where there are still an estimated 34 million current nicotine cigarette smokers[2]. Despite the known health risks, most quit attempts fail within a week[3] even with the aid of pharmacological and/or behavioral interventions[4]. A primary obstacle to quitting is the generally negative phenomenology of the Nicotine Withdrawal Syndrome (NWS) brought on following nicotine’s acute absence, with an aversive constellation of somatic, cognitive and affective symptoms peaking in the immediate days following abstinence. The severity of an individual’s dependence is therefore driven largely by an inability to withstand such periods of non-use. Mounting evidence from human neuroimaging studies indicates that much of the NWS phenomenology that arises, including craving, can be traced to alterations in specific brain circuits and networks [5-10]. As such, advancing the understanding of how brain circuitry reconfigures during this critical time window, how such abstinent “state” misconfigurations interact with underlying nicotine dependence “trait” misconfigurations associated with long-term nicotine use, and whether certain misconfigurations are related specifically to withdrawal severity, has the potential to drive more efficacious circuit-guided treatment approaches.

Among the most extensively implicated circuits in the pathology of nicotine dependence and SUD more broadly is the corticostriatal system. This circuitry is characterized by monosynaptic projections that transmit information processed within the frontal cortex to the striatum[11], the major input nucleus to the basal ganglia. The terminal fields of these projections form a topographic organization, wherein ventral and medial cortical regions project preferentially to ventromedial striatum (i.e., nucleus accumbens and ventral caudate and putamen) and dorsal and lateral cortical regions project preferentially to dorsolateral striatum (i.e., central and dorsal caudate and putamen)[12]. This projection topography influences the distinct functions and roles in SUD pathology played by striatal gross anatomical subdivisions (i.e., caudate, putamen, accumbens) and their frontal cortical connections. For instance, the nucleus accumbens and its frontal cortical connections have established roles in motivational and affective processes and are implicated in aberrant reward processing and craving in SUD[13-15]. The caudate nucleus and its frontal cortical connections subserve cognitive functions central to goal-directed behavior, while the putamen and its frontal cortical connections are implicated in the execution of stimulus-response habit behaviors[16, 17]. Transfer of the control of drug-use behavior from goal-directed processes mediated within the caudate to habit processes mediated within the putamen is thought to be a central mechanism underlying the development of compulsive drug use [18, 19].

One of the most ubiquitous tools used to noninvasively study corticostriatal circuits in SUD is resting-state functional connectivity (rsFC), which measures the strength of spatiotemporal coherence patterns of spontaneous neural activity (manifest as a BOLD hemodynamic signal) between different brain areas or networks[20]. Nearly two decades of work has found evidence that the effects of acute and chronic nicotine use in both nicotine sated and short-term abstinent smokers manifest as dysregulated functional connectivity in striatal cortical circuits[7, 10, 21-23]. The use of rsFC to investigate how this circuitry becomes aberrant in SUD commonly focuses on evaluating discrete A-to-B connections (i.e., between cortex-striatum node-pairs), and examining how their properties differ between groups (e.g., smokers vs. non-smokers) or states (e.g., drug satiety vs. acute abstinence), correlate with symptom severity (e.g., scores on the Fagerstrom Test for Nicotine Dependence) or predict treatment outcomes[24] [25]. Yet, emerging insights from quantitative neuroanatomy/tract-tracing in non-human primates underscore several properties of this circuitry’s connectional architecture that motivate the development of more sophisticated methods for evaluating its connectivity.

For example, anterograde tracing data have demonstrated that striatal terminal fields from different cortical regions overlap within the broader striatal topographic gradient, such that individual striatal nodes receive input from numerous cortical regions[26, 27]. Furthermore, retrograde tracing data that has quantified the relative strengths of the connections comprising striatal nodes’ “connectivity profiles”[28] has cast clearer light on 1) the comparatively modest input that any given cortical region contributes to a striatal node, and 2) the large number of cortical regions that contribute input of non-negligible strength to a given striatal node[29-31]. For instance, across nine nodes examined in different regions of the macaque striatum[29, 30], the strongest frontal cortical input region accounted for, on average, only 27.6% of the total input a striatal node received, with the remaining input contributed by connections from an average of 14.7 regions that each contributed more than 1% of the total. This distributed connectional structure has been observed in cortico-cortical circuits as well: across eight nodes along a rostro-caudal gradient of the macaque anterior cingulate cortex[31], each node’s strongest input region accounted for no more than 30% of its total anatomical input, with an average of 5.8 different regions accounting for 75% of a node’s total afferent connections. While other factors affect the relative influence of afferents on nodes (e.g., where they synapse on the cell, their temporal firing patterns), a direct relationship has been demonstrated between input cell quantity and functional connectivity strength[32-34]. Moreover, the electrophysiological properties of striatal projection neurons dictate that individual cortical inputs are insufficient for driving activation, and that only the spatiotemporally coordinated activity of multiple cortical inputs can drive striatal neurons to a sufficiently depolarized state where action potentials can occur [11, 35, 36].

Thus, if the net activity of brain nodes is shaped more by combinational features of multivariate connectivity profiles than by any individual afferent, methods designed to detect changes and aberrance in a node’s connectivity profile may provide important insights about the kinds and locations of circuit misconfigurations that are most central to the etiology and maintenance of neuropsychiatric disorders, including SUDs like nicotine dependence. As such, we introduce a connectivity profile analysis (CPA) approach for quantifying between- and within-group differences in voxel-wise, inter-regional connectivity profiles and identify sites of statistically significant difference using null modelling[37]. Since a node’s connectivity profile could be altered along several different dimensions – each of which may reflect a distinct form of circuit plasticity – we begin by defining and computing three complementary metrics for measuring connectivity profile misconfiguration: aggregate divergence, rank order misarrangement, and entropy shift. Using these metrics, we assess how voxel-wise connectivity profiles 1) differ tonically between non-smokers and smokers and 2) change phasically between a nicotine sated and acute abstinence state, when withdrawal symptoms often lead to relapse. Within smokers we also examine how interindividual differences in connectivity profile properties relate to dependence severity and withdrawal symptomology. Given the prominent role of aberrant corticostriatal circuits in both SUDs in general[19, 38, 39] and in nicotine dependence in particular[10, 21, 40-43], we focus this initial application of the CPA method on striatal connectivity profiles with the frontal cortex.

## Materials and Methods

### Participants

#### Empirical Sample

Participants were right-handed, aged 18–55 years, free of active drug or alcohol abuse/dependence (other than nicotine dependence in smokers), reporting no current psychiatric or neurological disorders, and presenting no contraindications for magnetic resonance imaging (MRI). Fifty-four current smokers and 35 non-smoking controls completed all experimental procedures. Following preprocessing, data from 8 smokers and 2 non-smokers were excluded due to excessive head motion (see below). Therefore, the final empirical sample for analysis consisted of 46 smokers (27 male; mean age 38.20 ± 11.99, mean years of education 13.00 ± 1.92) and 33 non-smokers (21 male; mean age 36.48 ± 9.57, mean years of education 13.55 ± 1.95). Groups did not differ in age, t(77)=0.719, p=0.474, sex, *X*^2^(1,79)=0.197, p=0.657, or years of education, t(77)=1.236, p=0.220. Written informed consent was obtained in accordance with the National Institute on Drug Abuse, Intramural Research Program Institutional Review Board.

### Normative Sample

The CPA procedure requires the computation of a normative distribution for each connectivity profile metric, to facilitate statistical assessment of empirically observed values. To do so, a normative sample from the Human Connectome Project (HCP) was curated to match the empirical sample (i.e., sample size, demographics, head motion) except for the presence of the group difference of interest in the empirical sample (i.e., smoking status). This allows the observation of empirically measured values that fall significantly outside the distribution of normative values to be attributed to the group difference of interest in the empirical sample. Procedures used to curate this matched normative sample are described in the **Supplemental Methods**.

### Experimental Design

Each non-smoker completed one MRI scanning session, and each smoker completed two MRI scanning sessions: one immediately following *ad lib* smoking, and the second during acute abstinence ∼48 hours after smoking the last cigarette prior to scanning. Since these data were part of an extended treatment protocol, the order of the two scans was fixed, with the sated scan preceding the abstinent scan by an average of 67 days (median 28 days). Further details on biochemical verification of abstinence and on subject exclusion criteria are provided in the **Supplemental Materials**.

### MRI Data Acquisition

Whole-brain echo planar images for 57 subjects (26 non-smokers and 31 smokers) were acquired on a 3T Siemens Trio scanner (Erlangen, Germany) using a 12-channel head coil. For resting-state data, 39 oblique axial slices (4-mm thick; 30° to anterior commissure–posterior commissure line) were acquired using a T2*-weighted, single-shot gradient echo, echo planar imaging sequence sensitive to blood oxygenation level-dependent (BOLD) effects (241 volumes; repetition time (TR)=2000□ ms; echo time (TE)=27□ms; flip angle (FA)=78°; field of view 220 × 220□mm^2^; image matrix 64 × 64). High-resolution oblique–axial T1-weighted structural images were acquired using a 3D magnetization-prepared rapid gradient-echo (MPRAGE) sequence (TR=1900□ms; TE=3.51□ms; TI=900□ms; FA=9°; voxel size=1□mm^3^). Images for 22 subjects (7 non-smokers and 15 smokers) were acquired on a Prisma using a 20-channel head coil and the same acquisition parameters as above.

The proportion of subjects scanned on each scanner did not differ between the non-smoker and smoker groups, *X*^2^(1,79)=1.24, p=0.265. Furthermore, after preprocessing with the same pipeline (see below), no significant differences in connectivity Z-score maps were observed across subjects in relation to which scanner was used. As such, images from both scanners were analyzed together.

### Resting-State fMRI Preprocessing

Preprocessing was performed using FMRIPREP version 20.2.1. A detailed description of these preprocessing steps is included in the **Supplemental Methods**. Further preprocessing included spatial blurring with a 6-mm full-width half-maximum Gaussian kernel and temporal filtering (0.01<f <0.1 Hz). To control for subject head motion, volumes were censored for framewise motion displacement (i.e., volume to volume movement) [44, 45]. The following frames were censored: frames with a framewise displacement (FD) > 0.5mm, frames preceding those with a FD > 0.5mm, and the first three frames of each scan. Subjects with more than 25% of frames censored were excluded from analysis. Eight smokers and two non-smokers were excluded from final analyses due to these head motion criteria.

Regions of interest (ROIs) used for generation of voxel-wise striatal fingerprints were based on the frontal cortical parcellation definitions from the Harvard-Oxford cortical atlas (**Supplementary Figure 1**), and included: supplementary motor cortex, superior frontal gyrus, subcallosal cortex, precentral gyrus, paracingulate gyrus, middle frontal gyrus, insular cortex, inferior frontal gyrus pars opercularis, inferior frontal gyrus pars triangularis, frontal pole, frontal orbital cortex, frontal operculum cortex, frontal medial cortex, central opercular cortex, and anterior cingulate cortex. The right hemisphere and left hemisphere component of each ROI was parcellated separately. To create sample-specific right and left striatum masks, the union of each subject’s caudate, putamen, and nucleus accumbens FreeSurfer parcellations was taken (separately for each hemisphere), and binary masks were created from the overlap of voxels present in at least 80% of subjects.

Resting-state functional connectivity (rsFC) between each striatal voxel and each frontal cortical seed (ROI) was assessed using the mean resting-state BOLD time series from each ROI extracted from each participant, which was then included in a GLM with 17 additional regressors of no interest: six motion parameters (three translations and three rotations) obtained from the rigid-body alignment of EPI volumes and their six temporal derivatives; the mean time series extracted from white matter; the mean times series extracted from CSF; and a second-order polynomial to model baseline signal and slow drift. The output of *r* values from the GLM were converted to Z-scores using Fisher’s *r*-to-Z transformation. Finally, given group differences in head-motion (**Supplementary Figure 2**), framewise displacement-adjusted Z-score maps were computed and served as the inputs to all main analyses (see **Supplementary Methods** for details and procedure).

### Resting-State fMRI Analysis

As the foundation for our analytic pipeline, we first established a frontal cortex connectivity profile (“fingerprint”)[46] for each voxel in the striatum. Each fingerprint quantified the strength of rsFC between a given striatal voxel and 30 (15 ipsilateral, 15 contralateral) frontal cortical subregions (‘targets’) [47], and was encoded by 30 voxel-wise striatal Z-score maps (one for each ROI) for each subject. Subject level fingerprints were then used to create three sets of group-level fingerprints – one averaged set each for sated smokers, abstinent smokers, and non-smokers.

Subsequently, we established three complementary methods for quantifying differences between voxels’ connectivity profile fingerprints across groups: aggregate divergence, rank order misarrangement, and entropy shift (**Supplementary Figure 3**). Aggregate divergence, quantified via Manhattan distance[46], measures the absolute cumulative magnitude by which all matched connections across two connectivity profiles differ. Rank order misarrangement measures how the order of strongest to weakest connections differs between two connectivity profiles. Entropy shift measures differences in the degree to which overall connectivity profile strength is concentrated in a few connections or distributed appreciably across many connections.

### Aggregate Divergence

To measure aggregate divergence between the connectivity profile of smokers and non-smokers at each voxel in the striatum, we carried out the following procedure [47] (**Supplementary Figure 4**) to separately compare the average striatal connectivity profiles of non-smokers to 1) the average striatal connectivity profiles of smokers during a nicotine sated state and 2) the average striatal connectivity profiles of smokers following a 48-hour abstinent state.

First, we computed voxel-wise striatal Z-score difference maps by taking the absolute value of the difference between the average smoker Z-score map and the average non-smoker Z-score map for each ROI (separately for the sated and abstinent states) – e.g.:

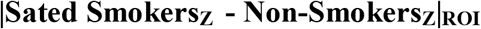

This produced 30 ROI difference maps for each smoking state. Then, for each state, we took the sum of the 30 ROI difference maps, resulting in final striatal voxel-wise maps whose values represent the Manhattan distance[46] between corticostriatal fingerprints of the average non-smoker and 1) the average sated smoker and 2) the average abstinent smoker – e.g.:

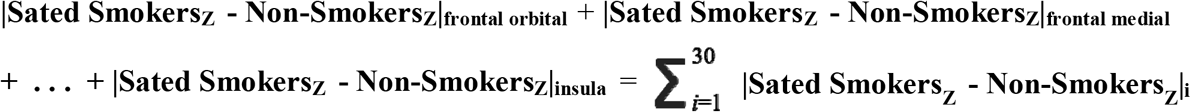

### Rank Order Misarrangement

Instead of constructing striatal voxel fingerprints using the Z-score connectivity values with each frontal cortical ROI, here we used “rank order fingerprints” (**Supplementary Figure 3b**) in which the value of each target ROI was its connectivity strength rank relative to the other ROIs in the fingerprint. As above, this procedure was carried out separately to compare non-smokers to 1) smokers during a nicotine sated state and 2) smokers during a 48-hour abstinent state.

After computing the rank order of each of the 30 cortical ROIs at each striatal voxel in each group based on the group-average ROI Z-score maps, we computed the absolute value of the difference between the rank order of each ROI at each voxel in the smoker group and non-smoker group. The sum of these 30 difference scores was then calculated to represent the “rank order misarrangement” between the groups at each voxel – e.g.:

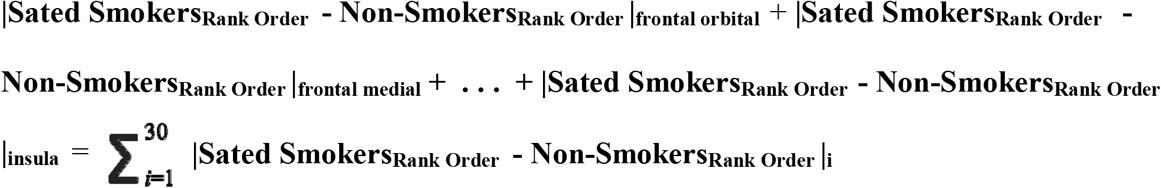

### Entropy Shift

Using the group-average Z-score connectivity voxel fingerprints, we calculated the entropy – a measure that indexes the uniformity of a distribution – of each voxel’s fingerprint in each group:

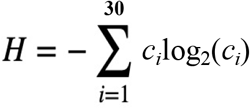

where *c*_*i*_ is the magnitude of connectivity with the *i-*th frontal cortical ROI[48]. For each voxel we divided this measure by log_2_30 to normalize it and bind it to the interval [0,1][49]. Higher values of entropy represent connectivity profiles where connectivity strength is more evenly distributed across ROIs. Conversely, lower values of entropy represent connectivity profiles where connectivity strength is more concentrated with one or a few ROIs. The difference in each group’s entropy at each voxel was than computed to represent the “entropy shift” between the groups (**Supplementary Figure 3c**).

### Determination of Significance Thresholds for Group-Level Analysis

To determine whether empirically observed voxel-level connectivity profile aberrance metrics (i.e., aggregate divergence, rank order misarrangement, and entropy shift) were statistically significant, we established a normative baseline of comparison using the matched “null” data from the HCP sample, which was also FD-adjusted in the same manner as the empirical sample. To do so, we first randomly split the HCP sample into two groups of sizes equal to those in the empirical sample (i.e., n=33 and n=46). We then calculated the aggregate divergence, rank order misarrangement, and entropy shift between the group average connectivity fingerprints at each striatal voxel in the same manner as for the empirical sample. After storing the resulting distributions, group membership was randomly permuted, and the process was repeated 10,000 times. The data from each permutation was combined to establish a final normative distribution for each of the three metrics. Voxel-wise p<0.001 thresholds for each metric were determined as: aggregate divergence ≥ 2.675; rank order misarrangement ≥ 160; entropy shift ≥ 0.045 (**Supplementary Figure 5a-c**). Corrected thresholding was set using 3dClustSim (AFNI 20.1.14) at p<0.001 uncorrected and k>7 to yield a p_FWE_ < 0.05.

### Within-Group Analysis

To identify striatal areas where connectivity profiles undergo significant within-smoker change following the transition from satiety to 48-hour abstinence, we first computed subject-level maps for each of the three misconfiguration metrics. This was done by replacing the smoker-average voxel-wise Z-score maps (as described above) with the individual subject maps and evaluating each smoker’s connectivity profiles (in each state) with respect to the non-smoker group-average connectivity profiles. Then, for each of the three metrics, we performed a voxel-wise paired t-test comparing the maps of smokers in the sated and 48-hour abstinent states.

#### Connectivity Profile Misconfiguration, Dependence Severity and Withdrawal Symptomology

Lastly, we sought to determine whether interindividual variance in smokers’ connectivity profile properties were associated with differences in dependency severity, as measured by the Fagerstrom Test for Nicotine Dependence (FTND)[50], and withdrawal symptomology during acute abstinence, as measured by the Wisconsin Withdrawal Scale (WSWS)[51]. Controlling for age, sex and years of education, we used linear regression to assess relationships between these metrics and subject-level misconfiguration metrics.

### Data availability

The HCP is an open-access project. HCP data used in this study can be downloaded by registering at https://db.humanconnectome.org. Anonymized empirical sample (i.e., smoker and non-smoker) data will be shared by request from qualified investigators. Custom scripts for the analyses are openly available at https://github.com/ckorponay/Connectivity-Profile-Misconfiguration.

## Results

### Aggregate Divergence

After two days of verified abstinence, large sections of the right and left medial and ventral striatum in smokers displayed significant aggregate divergence relative to non-smokers (**Figure 1a**). The sites of maximum aggregate divergence were in the right nucleus accumbens, bilateral caudal ventral putamen, and left dorsal caudate (**Table 1**). In contrast, we did not identify any evidence of significant aggregate divergence in these same smokers when in the nicotine sated state (**Figure 1b**).

**Figure 1:**
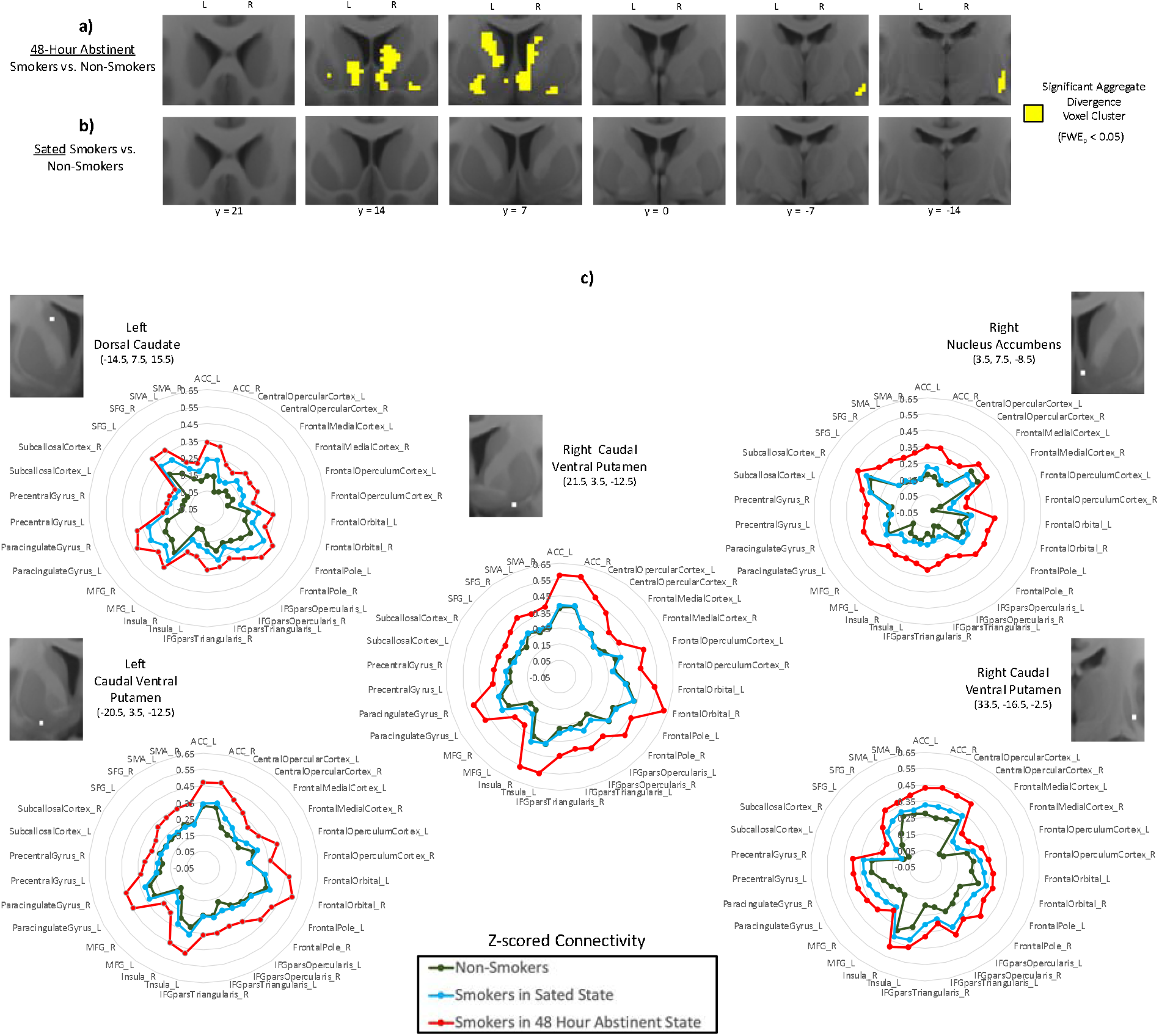
Aggregate divergence of striatal connectivity profiles in smokers relative to non-smokers. a) Significant aggregate divergence (yellow) emerges in the medial and ventral striatum in smokers after 48 hours of verified abstinence, but b) no aggregate divergence is evident when smokers are in a nicotine-sated state. c) The connectivity profiles (“fingerprints”) of non-smokers (dark green), sated smokers (blue), and acutely abstinent smokers (red) at the striatal sites of peak aggregate divergence (Table 1) in acutely abstinent smokers. Distance of each point from center of plot denotes Z-scored functional connectivity strength between striatal site and labeled frontal cortical ROI. Gap between red line and green line illustrates aggregate divergence of the connectivity profile in acute abstinence.

**Table 1.** Sites of significant (pFWE < 0.05, cluster size > 7) aggregate divergence in acutely abstinent smokers relative to non-smokers.

| ***Groups*** | ***Cluster Region*** | ***Cluster Size (voxels)*** | ***Peak Coordinate*** | ***Peak Manhattan Distance*** |
| --- | --- | --- | --- | --- |
| Abstinent Smokers vs. Non-smokers | Right rostral medial striatum | 206 | (3.5, 7.5, -8.5) | 4.65 |
|  | Right caudal lateral striatum | 51 | (33.5, -16.5, -2.5) | 4.27 |
|  | Right caudal lateral striatum | 41 | (21.5, 3.5, -12.5) | 4.45 |
|  | Left rostral medial striatum | 163 | (-14.5, 7.5, 15.5) | 4.18 |
|  | Left caudal lateral striatum | 69 | (-20.5, 3.5, -12.5) | 4.00 |

Examination of the FC fingerprint of each group at each of the five peak coordinates (polar plots in **Figure 1c**) revealed that striatal FC with all frontal cortical ROIs was larger in abstinent smokers relative to both sated smokers and non-smokers. Within smokers, a significant increase in aggregate divergence following the transition from nicotine satiety to 48-hour abstinence was observed only in the left nucleus accumbens (k=11, peak-level t = 3.75 at -6, 6, -12) (**Supplementary Figure 6**).

### Rank Order Misarrangement

Significant rank order misarrangement was identified in the right and left dorsal and lateral striatum of smokers in both the acutely abstinent (**Figure 2a**) and nicotine sated state (**Figure 2b)**. 69.7% of voxels that displayed significant rank order misarrangement during acute abstinence also displayed significant rank order misarrangement during satiety (**Supplementary Figure 7**), underscoring the constancy of rank order misarrangement in smokers across states. Moreover, within smokers, no significant changes in rank order arrangement were identified following the transition from satiety to 48-hour abstinence, further supporting the “trait” nature of this configuration difference. Sites of maximum rank order misarrangement during both satiety and abstinence included the right rostral dorsal putamen, right caudal ventral putamen, and left caudal dorsal putamen (**Table 2**).

**Figure 2:**
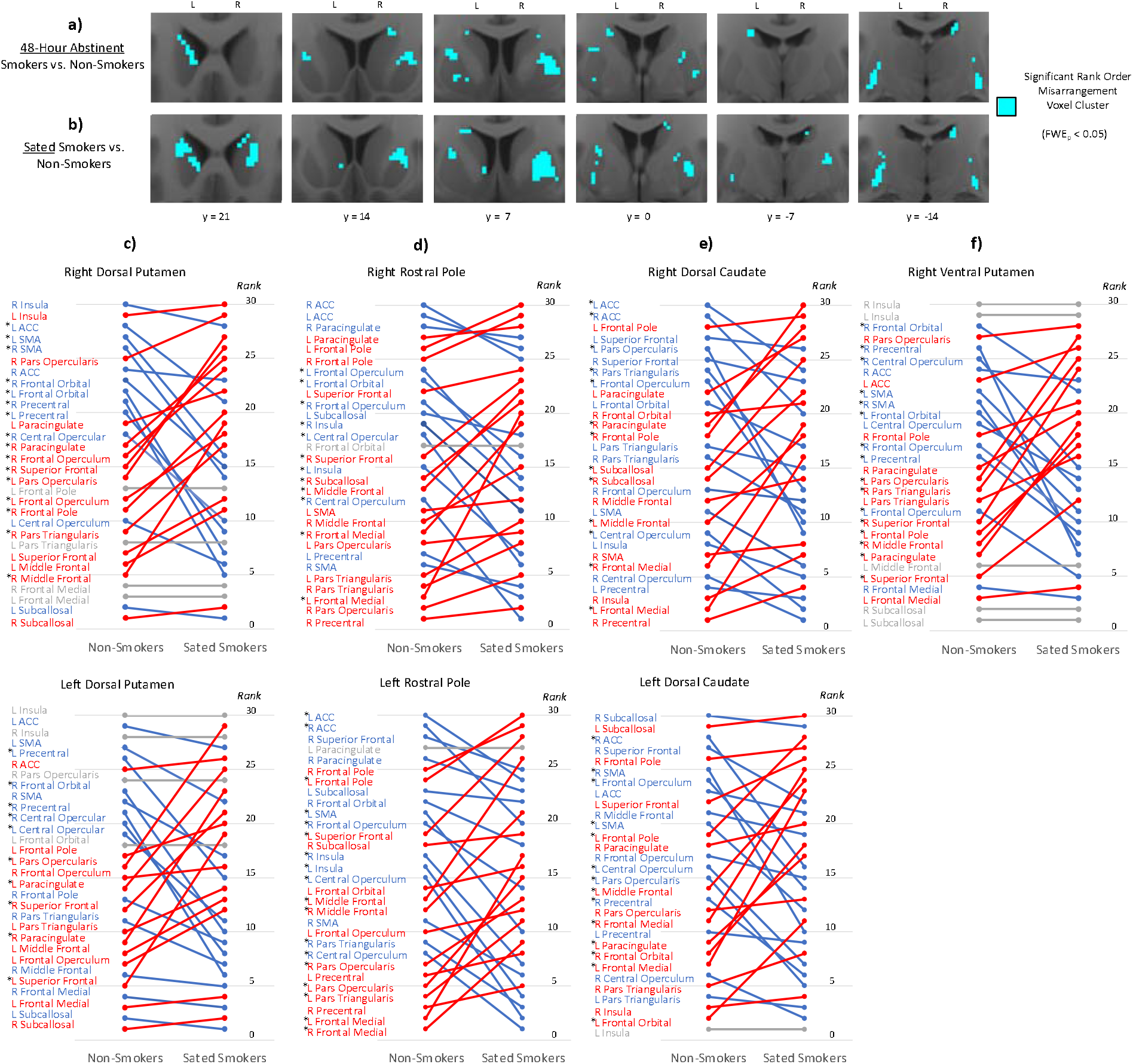
Rank order rearrangement of striatal connectivity profiles in smokers relative to non-smokers. Significant rank order rearrangement relative to non-smokers is present in smokers in both the a) acutely abstinent state and b) nicotine-sated state in the dorsal and lateral striatum (blue). c-f) At the striatal sites where rank order significantly differed between non-smokers and sated smokers (Table 2), illustrations of how the rank order arrangement of non-smokers (left) differed from that of sated smokers (right). Cortical ROIs at the top (i.e., with high ranks) indicate those with stronger connectivity with the striatal site, while cortical ROIs at the bottom (i.e., with low ranks) indicate those with weaker connectivity with the striatal site. Red indicates cortical ROIs whose rank was higher in sated smokers than in non-smokers, while blue indicates cortical ROIs whose rank was lower in sated smokers than in non-smokers. Gray indicates cortical ROIs whose rank was the same in both groups. Star denotes cortical ROIs whose rank order difference between non-smokers and sated smokers was statistically significant (p<0.001, rank order difference > 5).

**Table 2.** Sites of significant (pFWE < 0.05, cluster size > 7) rank order misarrangement in sated smokers and acutely abstinent smokers relative to non-smokers.

| *Groups* | *Cluster Region* | *Cluster Size (voxels)* | *Peak Coordinate* | *Peak Rank Order Difference* |
| --- | --- | --- | --- | --- |
| Sated Smokers vs. Non-smokers | Right rostral dorsal putamen | 238 | (23.5, 5.5, 5.5) | 232 |
|  | Right rostral pole | 66 | (19.5, 23.5, 5.5) | 282 |
|  | Right caudal dorsal caudate | 31 | (15.5, -14.5, 21.5) | 270 |
|  | Right caudal ventral putamen | 19 | (29.5, -14.5, -6.5) | 204 |
|  | Left caudal dorsal putamen | 154 | (-26.5, -12.5, 11.5) | 250 |
|  | Left rostral dorsal caudate | 54 | (-18.5, 19.5, 11.5) | 242 |
|  | Left rostral dorsal caudate | 12 | (-14.5, 5.5, 23.5) | 226 |
| Abstinent Smokers vs. Non-smokers | Right rostral dorsal putamen | 94 | (21.5, 11.5, 7.5) | 220 |
|  | Right caudal ventral putamen | 20 | (29.5, -18.5, -0.5) | 216 |
|  | Right rostral dorsal caudate | 19 | (17.5, 15.5, 17.5) | 216 |
|  | Right caudal dorsal putamen | 14 | (27.5, -10.5, 9.5) | 206 |
|  | Right caudal dorsal caudate | 14 | (15.5, -12.5, 23.5) | 214 |
|  | Right rostral pole | 11 | (17.5, 25.5, -2.5) | 220 |
|  | Right caudal dorsal putamen | 9 | (23.5, -2.5, 11.5) | 180 |
|  | Left caudal ventral putamen | 49 | (-30.5, -20.5, 1.5) | 238 |
|  | Left caudal dorsal putamen | 46 | (-24.5, 1.5, 13.5) | 246 |
|  | Left caudal dorsal caudate | 27 | (-18.5, -4.5, 25.5) | 208 |
|  | Left rostral ventral putamen | 19 | (-22.5, 5.5, -4.5) | 190 |
|  | Left rostral pole | 18 | (-10.5, 21.5, 3.5) | 218 |

Within both the right and left dorsal putamen clusters, connectivity strength rank was significantly lower in sated smokers compared to non-smokers for right precentral gyrus (5 versus 21 in right striatum, 6 versus 21 in left striatum), left precentral gyrus (9 versus 20 in right striatum, 15 versus 26 in left striatum), and right frontal orbital cortex (15 versus 23 in right striatum, 10 versus 23 in left striatum) (**Figure 2c**). Conversely, rank was significantly larger in sated smokers compared to non-smokers for right superior frontal gyrus (26 versus 15 in right striatum, 21 versus 12 in left striatum), and left inferior frontal gyrus pars opercularis (27 versus 14 in right striatum, 29 versus 16 in left striatum) (**Figure 2c**). Collectively, whereas dorsal putamen connectivity in non-smokers is stronger with motor and limbic frontal areas than with more cognitive frontal areas, rank order arrangement in sated smokers is swapped such that dorsal putamen connectivity is stronger with cognitive frontal areas than with motor and limbic frontal areas.

Within the right and left rostral pole clusters, connectivity strength rank was significantly greater in sated smokers compared to non-smokers for bilateral frontal medial cortex, but significantly lower in sated smokers for bilateral insula (**Figure 2d**). The right and left dorsal caudate clusters were both characterized by significantly greater connectivity strength rank in sated smokers for left frontal medial cortex and left middle frontal gyrus, and significantly lower rank in sated smokers for right ACC, left inferior frontal gyrus pars opercularis, and left frontal opercular cortex (**Figure 2e)**. The right ventral putamen cluster was characterized by significantly lower connectivity rank in sated smokers with bilateral motor and premotor cortices and significantly greater connectivity rank with bilateral inferior and superior frontal gyri (**Figure 2f)**.

### Entropy Shift

Evidence for significant entropy shift of striatal connectivity profiles in smokers compared to non-smokers was minimal (**Table 3**), with a handful of small clusters displaying entropy shift primarily in the acute abstinence state. Within smokers, entropy shift relative to non-smokers was significantly greater during acute abstinence than during satiety in the left rostral putamen (k=16, peak-level t = 4.20 at -16, 12, -4) and left caudal caudate (k=10, peak-level t = 4.16 at -10, 2, 18) (**Supplementary Figure 6**).

**Table 3.**
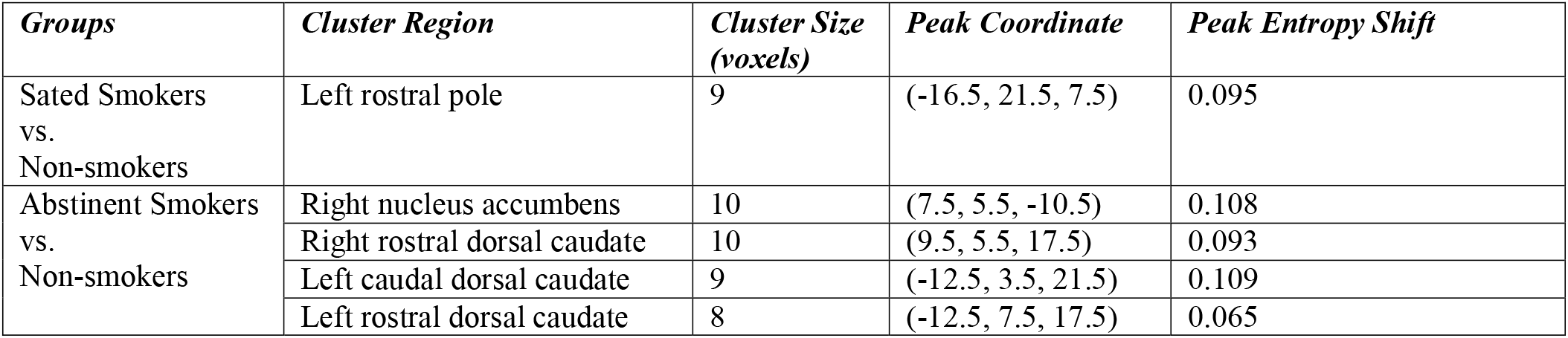
Sites of significant (pFWE < 0.05, cluster size > 7) entropy shift in sated smokers and acutely abstinent smokers relative to non-smokers.

| *Groups* | *Cluster Region* | *Cluster Size (voxels)* | *Peak Coordinate* | *Peak Entropy Shift* |
| --- | --- | --- | --- | --- |
| Sated Smokers vs. Non-smokers | Left rostral pole | 9 | (-16.5, 21.5, 7.5) | 0.095 |
| Abstinent Smokers vs. Non-smokers | Right nucleus accumbens | 10 | (7.5, 5.5, -10.5) | 0.108 |
|  | Right rostral dorsal caudate | 10 | (9.5, 5.5, 17.5) | 0.093 |
|  | Left caudal dorsal caudate | 9 | (-12.5, 3.5, 21.5) | 0.109 |
|  | Left rostral dorsal caudate | 8 | (-12.5, 7.5, 17.5) | 0.065 |

### Spatial Segregation and Overlap of Types of Connectivity Profile Misconfiguration

Striatal areas of significant aggregate divergence, rank order misarrangement, and entropy shift were largely non-overlapping (**Figure 3**). Of all identified voxels with significant aggregate divergence in the acute abstinence state, only 2.83% also displayed significant rank order misarrangement, and only 1.51% also displayed significant entropy shift. Likewise, of all identified voxels with significant rank order misarrangement in the acute abstinence state, only 4.41% also displayed significant aggregate divergence, and only 0.59% also displayed significant entropy shift. Spatially, this was reflected in a ventromedial/dorsolateral segregation, wherein significant aggregate divergence was primarily observed in medial and ventral striatum, while significant rank order misarrangement was primarily observed in lateral and dorsal striatum (**Figure 3**).

**Figure 3:**
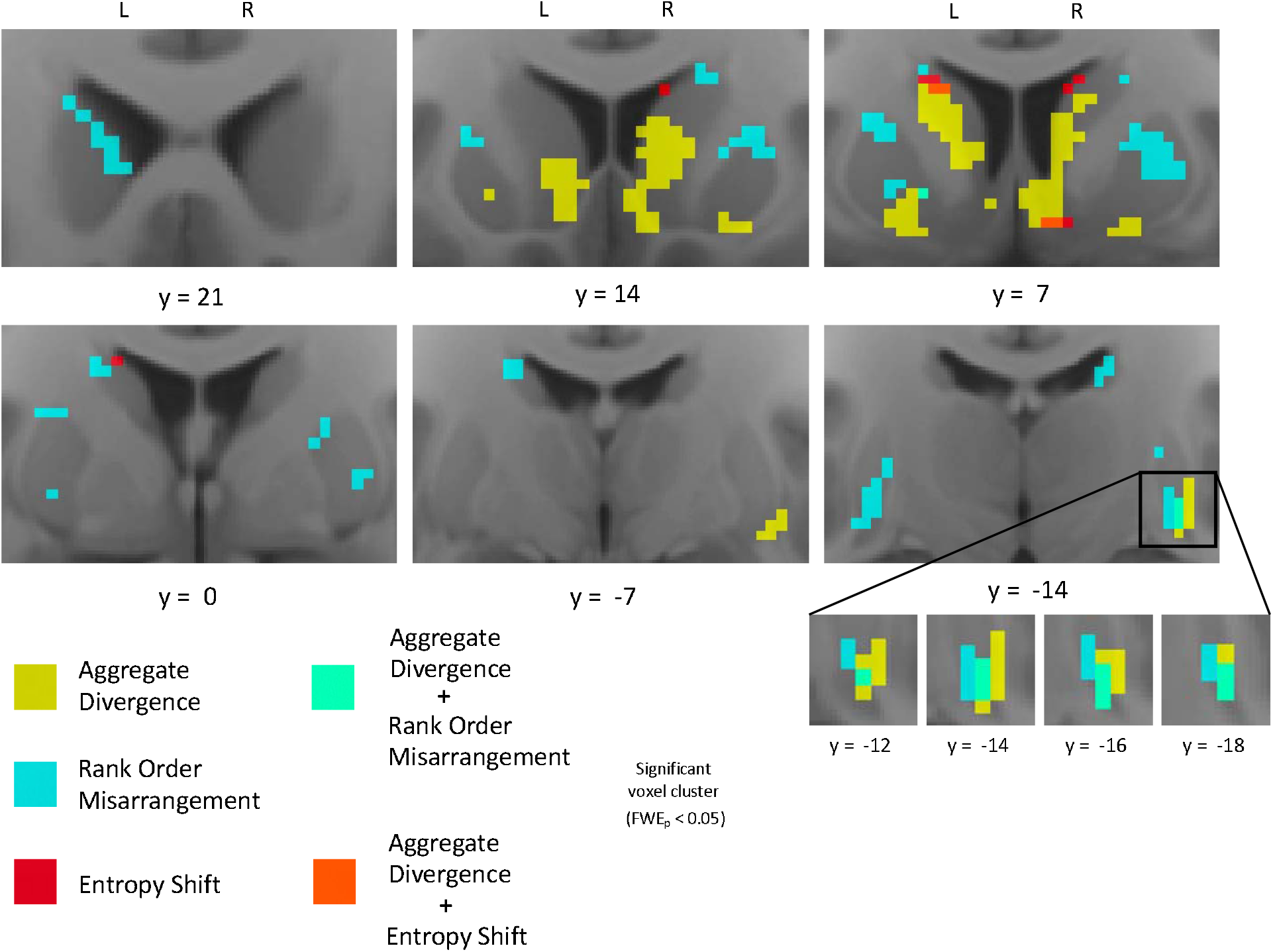
Spatial segregation of aggregate divergence and rank order rearrangement and their unique co-occurrence in the right caudal ventral putamen after 48-hour abstinence.

However, a cluster of size k=9 in right caudal ventral putamen uniquely displayed both significant aggregate divergence and significant rank order misarrangement in acutely abstinent smokers (**Figure 3**). Moreover, six contiguous voxels within this cluster also displayed significant rank order misarrangement in sated smokers. We explored this unique k=6 cluster in depth to determine precisely how these multiple types of connectivity profile misconfiguration manifested (**Figure 4**). In smokers during satiety, the connectivity strength rank of motor and premotor control areas in the ventral putamen is significantly lower compared to non-smokers, while the rank of cognitive processing areas is significantly higher. This results in a right caudal ventral putamen connectivity profile in sated smokers where the rank order of motor and cognitive frontal area connectivity strengths is swapped compared to non-smokers (**Figure 4a**). This swapped rank order arrangement is maintained in smokers following the transition from satiety to 48-hour abstinence (**Figure 4b**). However, though similar to non-smokers in aggregate connectivity magnitude during satiety (**Figure 4c**), this rearranged connectivity profile also becomes significantly greater in aggregate strength during acute abstinence (**Figure 4d**). This is driven by the emergence of significantly greater connectivity of right caudal ventral putamen with cognitive frontal areas – but not with motor or premotor frontal areas – compared to non-smokers.

**Figure 4:**
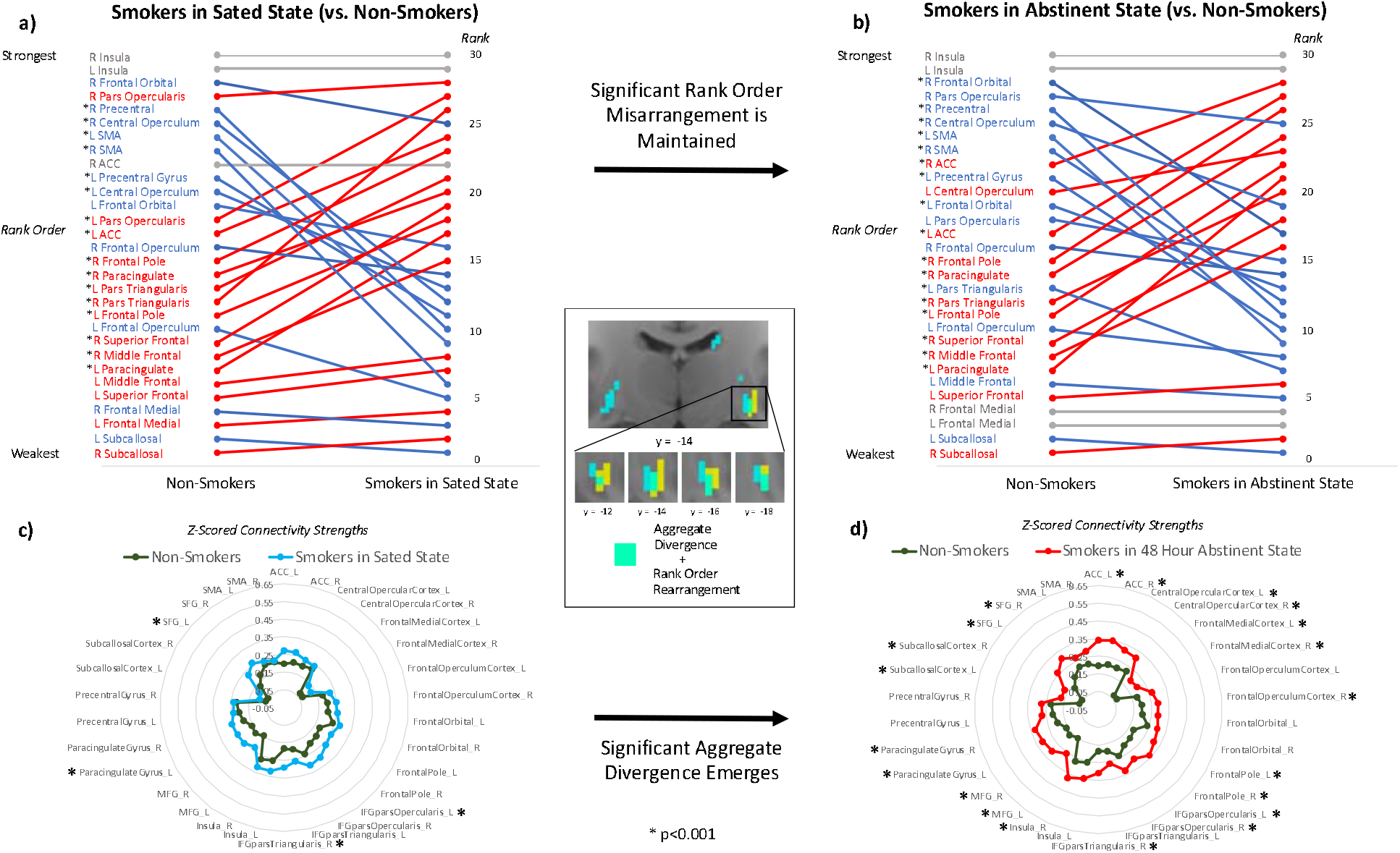
**Multiple forms of connectivity profile reconfiguration in the right caudal ventral putamen** (turquois voxels in center panel). Significant rank order rearrangement in smokers relative to non-smokers is evident during a) nicotine satiety, and is maintained following b) 48-hour abstinence. Significant aggregate divergence in smokers relative to non-smokers is not evident during c) nicotine satiety, but emerges after d) 48-hour abstinence.

### Connectivity Profile Misconfiguration, Dependence Severity and Withdrawal Symptomology

We next explored whether the interactive magnitude of aggregate divergence and rank order misarrangement in the k=6 right caudal ventral putamen cluster following the transition from satiety to acute abstinence was related to the severity of nicotine dependence and withdrawal symptomology across smokers. Controlling for age, gender, and education, as well as the main effects of aggregate divergence and rank order misarrangement, the interactive magnitude of these two connectivity profile misconfigurations was significantly negatively associated with dependence severity indexed by the FTND, t(37) = -2.510, p = 0.017. Low magnitudes of both misconfiguration types (i.e., connectivity profiles resembling non-smokers) and high magnitudes of both misconfiguration types predicted low FTND scores (**Supplementary Figure 8**). Conversely, a high magnitude of one misconfiguration type but a low magnitude of the other predicted high FTND scores. This was the only factor in the model significantly associated with dependence severity other than age, t(37) = 3.555, p=0.001. Notably, separate tests of the main effect of aggregate divergence, t(39)=0.480, p=0.634, and of rank order misarrangement, t(39)=-0.182, p=0.856, were not significant.

In contrast, these right caudal ventral putamen connectivity profile misconfigurations were not significantly associated with withdrawal symptomology during acute abstinence as indexed by the WSWS. However, given prior work implicating nucleus accumbens activity in nicotine withdrawal[52], we examined withdrawal severity in relation to aggregate divergence in the k=11 cluster identified in left nucleus accumbens where this misconfiguration metric increased significantly within smokers between satiety and acute abstinence. We found that, across smokers, greater levels of left nucleus accumbens aggregate divergence during acute abstinence were associated with greater withdrawal symptomology as indexed by WSWS total score during acute abstinence, t(39)=2.209, p=0.033, controlling for age, sex, and education (**Supplementary Figure 9**). Left nucleus accumbens aggregate divergence in acute abstinence was not significantly related to FTND dependence severity, t(39)=0.268, p=0.790.

FTND scores of dependence severity and WSWS scores of withdrawal severity were not significantly related to each other, t(39)=-0.003, p=0.998 (consistent with prior findings of weak univariate relationships between dependence severity and withdrawal[53-55]) nor were the interactive magnitude of aggregate divergence and rank order misarrangement in right caudal ventral putamen and aggregate divergence in left nucleus accumbens, t(37)=0.664, p=0.510.

## Discussion

Recent insights from neuroanatomical track tracing studies warrant increased examination of the integrity of nodal “connectivity profiles” – the multivariate set of connection strengths that brain nodes have with other areas of the brain [28, 56] – to better understand brain processes in both the healthy individual as well as in those with neuropsychiatric disorders like SUD. Yet, existing FC analysis approaches are not optimized for the identification of such “connectivity profile misconfigurations.” For instance, traditional univariate seed-based approaches (i.e., node-to-node, node-to-network, network-to-network, etc.) are effective detectors of aberrant individual connections, but do not gauge aggregated abnormality across connectivity profiles. Moreover, existing approaches that incorporate multivariate information about nodes’ connectivity profiles measure only ultra-local connections (e.g. regional homogeneity [57]), or, in the case of graph-theoretic approaches[58, 59], focus on abstracted network properties that may obscure important variance in underlying connection strengths.

Here, we introduce a connectivity profile analysis (CPA) approach for quantifying and statistically evaluating distinct types of connectivity profile misconfigurations and apply it to examine striatal connectivity profiles with frontal cortex in nicotine dependent smokers. First, we found evidence of significant smoking-related connectivity profile misconfigurations in largely bilateral areas of both the dorsal and ventral striatum. Foremost, these results suggest that prior findings of abnormal connections between individual pairs of striatal and cortical nodes[10, 21, 39] appear to represent the “tip of the iceberg” of corticostriatal circuit alteration in nicotine dependence. More specifically, they suggest that abnormal activity and function of striatal nodes in nicotine dependence may be related not simply to large changes in the strength of individual frontal connections, but to the accumulation of changes in frontal connections throughout connectivity profiles that in aggregate are substantial.

We further demonstrate that connectivity profile misconfiguration manifests in several different and largely independent forms, as indexed by the measures of aggregate divergence, rank order misarrangement, and entropy shift. Fewer than 5% of all striatal voxels that displayed significant connectivity profile misconfiguration in one form also displayed significant misconfiguration in another form. These findings support the idea of different types of connectivity profile misconfiguration that may each reflect a distinct form of neurobiological plasticity, and which may relate to a separate alteration in the node’s activity and function. For example, appreciable increases (or decreases) to connection strengths throughout the connectivity profile – indexed by aggregate divergence – may upregulate (or downregulate) the node’s responsiveness to marginal input and its overall level of activity[60, 61]. Separately, the reshuffling of which connections are stronger and weaker – indexed by rank order misarrangement – may adjust which regions have more or less influence in shaping the node’s activity[62-64] and thus change the input combinations that drive nodal activation and subsequent behavior. Moreover, alterations in the extent to which overall connection strength is concentrated in a few connections versus distributed evenly across many[31] – indexed by entropy shift – may change the relative magnitudes by which inputs influence nodal activity independent of changes to the rank order of their influence.

Notably, therefore, we found that bilateral dorsal and ventral striatum in smokers display distinct forms of connectivity profile misconfiguration. Aggregate divergence in smokers – which presented only following acute abstinence – was observed almost exclusively in the bilateral ventral striatum (i.e., nucleus accumbens and ventromedial caudate) and was characterized by a global, proportional increase in FC with all subregions of the frontal cortex without significant alterations to connectivity profile rank order or entropy. The nature of this connectivity profile misconfiguration suggests that acute abstinence increases the response of the ventral striatum to marginal frontal input and its overall level of activity, without altering the balance of how much each of its connections shapes its activity. Furthermore, we demonstrate that across smokers, higher levels of left nucleus accumbens aggregate divergence following the transition to acute abstinence were associated with greater withdrawal symptomology. The direction of these results is consistent with those from prior work that has found FC increases node-to-node, node-to-network, and network-to-network connections in smokers during short-term abstinence[7, 8, 21, 65, 66], which have also been associated with increases in subjective ratings of withdrawal[8, 67] and worse treatment outcomes[67]. For example, Janes and colleagues[8] report increases in FC between the medial and orbital prefrontal cortex and a range of brain regions including the striatum after just 1 hour of abstinence that are associated with increases in subjective craving. Here, we extend these findings by demonstrating that these changes are associated with alterations not just in A-to-B connections but more broadly in connectivity profile alterations.

In the bilateral dorsal striatum (i.e., putamen and dorsolateral caudate), on the other hand, connectivity profile misconfiguration in smokers was characterized almost exclusively by rank order misarrangement – and was observable during both acute abstinence and satiety relative to non-smokers. In non-smokers, putamen connectivity with motor (i.e., supplementary motor area, primary motor cortex) and limbic (i.e., orbitofrontal cortex) frontal cortical areas was stronger than that with cognitive frontal areas (i.e., dorsolateral prefrontal cortex, ventrolateral prefrontal cortex, frontal pole). This configuration was swapped in smokers (i.e., connections with cognitive frontal areas were stronger than those with motor and limbic frontal areas), and this misarrangement was maintained regardless of smoking state (i.e., sated or acutely abstinent). Rank order misarrangement in the caudate in smokers involved changes amongst different limbic and cognitive frontal areas. Here, the strength ranks of connections with frontal medial cortex and middle frontal gyrus are significantly higher relative to non-smokers, while those with anterior cingulate cortex, pars opercularis of the inferior frontal gyrus, and frontal opercular cortex are significantly lower relative to non-smokers. As in the ventral striatum, the predominant form of connectivity profile misconfiguration in the dorsal striatum (i.e., rank order misarrangement) largely occurred without the presence of other forms (i.e., aggregate divergence or entropy shift). This suggests that while dorsal striatum may be no more or less active or responsive to marginal input in smokers compared to non-smokers, the relative influence that individual frontal cortical areas have on shaping dorsal striatal activity are shuffled – for example, from more motor region influence to more cognitive region influence in the dorsal putamen. Moreover, the stability of these misarrangements across smoking states suggests that these circuit misconfigurations are “trait” properties of long-term smoking. Speculatively, these connectivity profile rank order misarrangements in the dorsal striatum could reflect alterations associated with the biasing of behavioral control from the goal-directed to the habitual behavior system that is posited to occur in addiction[19, 68]. While the neural basis for this shift has primarily focused on striatal interactions with the dopaminergic midbrain[18], the current findings provide a window for investigating the potential role of frontostriatal interactions in this change.

Notably, we identified one focal area of the striatum – right caudal ventral putamen – that uniquely displayed both trait rank order misarrangement and state-dependent aggregate divergence in smokers. Here, connections with cognitive frontal areas (i.e., ventrolateral prefrontal cortex, dorsolateral prefrontal cortex, frontal pole) became stronger than those with motor frontal areas (i.e., primary motor cortex and supplementary motor area) in the connectivity profile as a trait-level misconfiguration. Moreover, aggregate divergence during acute abstinence amplified the difference in strength between these classes of connections, as the increases in connection strengths with cognitive areas were statistically significant while those with motor/premotor areas were not. Furthermore, when examined across smokers, we found that the interactive magnitude of these two misconfiguration types at this striatal site was significantly associated with dependence severity (i.e., FTND), wherein low magnitudes of both misconfiguration types (i.e., connectivity profiles resembling non-smokers) and high magnitudes of both misconfiguration types predicted low FTND scores. Yet, there was no relationship with the severity of withdrawal symptomology (i.e., WSWS). Collectively, the findings suggest that these connectivity profile misconfigurations in the right caudal ventral putamen may serve a compensatory function that helps to attenuate dependence during nicotine absence, but via a mechanism other than reduction of aversive withdrawal symptoms. One potential mechanism could be an increase in inhibitory control over habit-driven behavior. This interpretation is consistent with findings that this region of the striatum has frequently been implicated in inhibitory control as well as habit learning and execution[69-72]. For example, a recent meta-analysis found that neural representations of both laboratory-produced habits and naturalistic habits acquired outside the laboratory are localized in the caudal putamen[70]. Misconfiguration of this component of the habit execution circuit during acute abstinence may serve to reduce the propensity for habit-driven smoking behavior despite negative withdrawal symptoms.

In sum, using a novel connectivity profile analysis approach, we identify nicotine dependence related connectivity profile misconfigurations in dorsolateral and ventromedial striatum that differ in both type and state-dependency, and identify a unique striatal site where the extent to which smoking trait misconfigurations are magnified by misconfigurations during the acute abstinence state is linked to dependence severity. Identified sites of maximal connectivity profile misconfiguration could potentially serve as a useful targeting guide for neuromodulation-based therapies and/or as a biomarker readout for treatment efficacy. Further research is warranted to investigate the potential linkages between connectivity profile misconfigurations and cognitive-behavioral functions, such as inhibitory control and reward-processing, as well as the potential significance of laterality effects. It will also be of interest to examine how neuromodulation affects misconfigured connectivity profiles and the degree to which it normalizes them, and how targeting sites identified via a connectivity profile analysis approach impacts treatment outcomes. Furthermore, in conjunction with a recently developed connectome-based predictive modeling “computational lesion” approach[73, 74], which identifies nodes whose connectivity profile contributes most to the predictive power of a model, the current connectivity profile-centered approach may also have potential for prediction of treatment outcome success as well as a marker of resilience to develop drug dependence.

## Acknowledgements

This work utilized the computational resources of the NIH HPC Biowulf cluster (http://hpc.nih.gov).

## References

1. Health, U.D.o. and H. Services, The health consequences of smoking—50 years of progress: a report of the Surgeon General. Atlanta (GA): Centers for Disease Control and Prevention. national Center for Chronic disease Prevention and Health Promotion, office on smoking and Health, 2014.

2. Creamer, M.R., et al., Tobacco product use and cessation indicators among adults— United States, 2018. Morbidity and mortality weekly report, 2019. 68(45): p. 1013.

3. Hughes, J.R., J. Keely, and S. Naud, Shape of the relapse curve and long-term abstinence among untreated smokers. Addiction, 2004. 99(1): p. 29–38.

4. Ohara, P., et al., Design and results of the initial intervention program for the Lung Health Study. Preventive medicine, 1993. 22(3): p. 304–315.

5. Aronson Fischell, S., et al., Transcranial Direct Current Stimulation Applied to the Dorsolateral and Ventromedial Prefrontal Cortices in Smokers Modifies Cognitive Circuits Implicated in the Nicotine Withdrawal Syndrome. Biol Psychiatry Cogn Neurosci Neuroimaging, 2020. 5(4): p. 448–460.

6. Sutherland, M.T., et al., Resting state functional connectivity in addiction: Lessons learned and a road ahead. Neuroimage, 2012. 62(4): p. 2281–95.

7. Fedota, J.R. and E.A. Stein, Resting-state functional connectivity and nicotine addiction: prospects for biomarker development. Ann N Y Acad Sci, 2015. 1349: p. 64–82.

8. Janes, A.C., et al., An increase in tobacco craving is associated with enhanced medial prefrontal cortex network coupling. PloS one, 2014. 9(2): p. e88228.

9. Lerman, C., et al., Large-scale brain network coupling predicts acute nicotine abstinence effects on craving and cognitive function. JAMA psychiatry, 2014. 71(5): p. 523–530.

10. Hong, L.E., et al., Association of nicotine addiction and nicotine’s actions with separate cingulate cortex functional circuits. Archives of general psychiatry, 2009. 66(4): p. 431–441.

11. Wilson, C., The contribution of cortical neurons to the firing pattern of striatal spiny neurons. Model of Information Processing in Basal Ganglia, 1995. 1: p. 29–31.

12. Haber, S.N., Corticostriatal circuitry. Dialogues in clinical neuroscience, 2016. 18(1): p. 7.

13. David, S.P., et al., Ventral striatum/nucleus accumbens activation to smoking-related pictorial cues in smokers and nonsmokers: a functional magnetic resonance imaging study. Biological psychiatry, 2005. 58(6): p. 488–494.

14. Koob, G.F. and M. Le Moal, Drug addiction, dysregulation of reward, and allostasis. Neuropsychopharmacology, 2001. 24(2): p. 97–129.

15. Robinson, T.E. and K.C. Berridge, The neural basis of drug craving: an incentive-sensitization theory of addiction. Brain research reviews, 1993. 18(3): p. 247–291.

16. Balleine, B.W. and J.P. O’doherty, Human and rodent homologies in action control: corticostriatal determinants of goal-directed and habitual action. Neuropsychopharmacology, 2010. 35(1): p. 48–69.

17. de Wit, S., et al., Corticostriatal connectivity underlies individual differences in the balance between habitual and goal-directed action control. Journal of Neuroscience, 2012. 32(35): p. 12066–12075.

18. Everitt, B.J. and T.W. Robbins, Neural systems of reinforcement for drug addiction: from actions to habits to compulsion. Nature neuroscience, 2005. 8(11): p. 1481–1489.

19. Everitt, B.J. and T.W. Robbins, Drug addiction: updating actions to habits to compulsions ten years on. Annual review of psychology, 2016. 67: p. 23–50.

20. Biswal, B., et al., Functional connectivity in the motor cortex of resting human brain using echo-planar MRI. Magnetic resonance in medicine, 1995. 34(4): p. 537–541.

21. Claus, E.D., et al., Association between nicotine dependence severity, BOLD response to smoking cues, and functional connectivity. Neuropsychopharmacology, 2013. 38(12): p. 2363–2372.

22. Brooks, S., J. Ipser, and D. Stein, Chronic and acute nicotine exposure versus placebo in smokers and nonsmokers: A systematic review of resting-state fmri studies. Addictive Substances and Neurological Disease, 2017: p. 319–338.

23. Scarlata, M.J., R.J. Keeley, and E.A. Stein, Nicotine addiction: Translational insights from circuit neuroscience. Pharmacol Biochem Behav, 2021. 204: p. 173171.

24. Sutherland, M.T., et al., Insula’s functional connectivity with ventromedial prefrontal cortex mediates the impact of trait alexithymia on state tobacco craving. Psychopharmacology, 2013. 228(1): p. 143–155.

25. Weiland, B.J., et al., Reduced executive and default network functional connectivity in cigarette smokers. Human brain mapping, 2015. 36(3): p. 872–882.

26. Averbeck, B.B., et al., Estimates of projection overlap and zones of convergence within frontal-striatal circuits. Journal of Neuroscience, 2014. 34(29): p. 9497–9505.

27. Haber, S.N., et al., Reward-related cortical inputs define a large striatal region in primates that interface with associative cortical connections, providing a substrate for incentive-based learning. Journal of Neuroscience, 2006. 26(32): p. 8368–8376.

28. Passingham, R.E., K.E. Stephan, and R. Kotter, The anatomical basis of functional localization in the cortex. Nat Rev Neurosci, 2002. 3(8): p. 606–16.

29. Korponay, C., E.Y. Choi, and S.N. Haber, Corticostriatal Projections of Macaque Area 44. Cerebral Cortex Communications, 2020. 1(1): p. tgaa079.

30. Choi, E.Y., S.-L. Ding, and S.N. Haber, Combinatorial inputs to the ventral striatum from the temporal cortex, frontal cortex, and amygdala: implications for segmenting the striatum. Eneuro, 2017. 4(6).

31. Tang, W., et al., A connectional hub in the rostral anterior cingulate cortex links areas of emotion and cognitive control. Elife, 2019. 8: p. e43761.

32. Hori, Y., et al., Comparison of resting-state functional connectivity in marmosets with tracer-based cellular connectivity. Neuroimage, 2020. 204: p. 116241.

33. Reid, A.T., et al., A cross-modal, cross-species comparison of connectivity measures in the primate brain. Neuroimage, 2016. 125: p. 311–331.

34. Miranda-Dominguez, O., et al., Bridging the gap between the human and macaque connectome: a quantitative comparison of global interspecies structure-function relationships and network topology. Journal of Neuroscience, 2014. 34(16): p. 5552–5563.

35. Wilson, C.J. and Y. Kawaguchi, The origins of two-state spontaneous membrane potential fluctuations of neostriatal spiny neurons. Journal of neuroscience, 1996. 16(7): p. 2397–2410.

36. Carter, A.G., G.J. Soler-Llavina, and B.L. Sabatini, Timing and location of synaptic inputs determine modes of subthreshold integration in striatal medium spiny neurons. Journal of Neuroscience, 2007. 27(33): p. 8967–8977.

37. Markello, R.D. and B. Misic, Comparing spatial null models for brain maps. NeuroImage, 2021. 236: p. 118052.

38. Kalivas, P.W., Addiction as a pathology in prefrontal cortical regulation of corticostriatal habit circuitry. Neurotoxicity research, 2008. 14(2): p. 185–189.

39. Hu, Y., et al., Compulsive drug use is associated with imbalance of orbitofrontal-and prelimbic-striatal circuits in punishment-resistant individuals. Proceedings of the National Academy of Sciences, 2019. 116(18): p. 9066–9071.

40. Xia, J., A.M. Meyers, and J.A. Beeler, Chronic nicotine alters corticostriatal plasticity in the striatopallidal pathway mediated by NR2B-containing silent synapses. Neuropsychopharmacology, 2017. 42(12): p. 2314–2324.

41. Janes, A.C., et al., Insula–dorsal anterior cingulate cortex coupling is associated with enhanced brain reactivity to smoking cues. Neuropsychopharmacology, 2015. 40(7): p. 1561–1568.

42. Keeley, R.J., et al., Intrinsic differences in insular circuits moderate the negative association between nicotine dependence and cingulate-striatal connectivity strength. Neuropsychopharmacology, 2020. 45(6): p. 1042–1049.

43. Hsu, L.-M., et al., Intrinsic insular-frontal networks predict future nicotine dependence severity. Journal of Neuroscience, 2019. 39(25): p. 5028–5037.

44. Power, J.D., et al., Spurious but systematic correlations in functional connectivity MRI networks arise from subject motion. Neuroimage, 2012. 59(3): p. 2142–54.

45. Yan, C.G., et al., A comprehensive assessment of regional variation in the impact of head micromovements on functional connectomics. Neuroimage, 2013. 76: p. 183–201.

46. Mars, R.B., et al., Comparing brains by matching connectivity profiles. Neuroscience & Biobehavioral Reviews, 2016. 60: p. 90–97.

47. Korponay, C., E.A. Stein, and T. Ross, Connectivity profile laterality in corticostriatal functional circuitry: a fingerprinting approach. bioRxiv, 2021.

48. Conrad, K., Probability distributions and maximum entropy. Entropy, 2004. 6(452): p. 10.

49. Faskowitz, J., et al., Edge-centric functional network representations of human cerebral cortex reveal overlapping system-level architecture. Nat Neurosci, 2020. 23(12): p. 1644–1654.

50. Dijkstra, A. and D. Tromp, Is the FTND a measure of physical as well as psychological tobacco dependence? Journal of substance abuse treatment, 2002. 23(4): p. 367–374.

51. Welsch, S.K., et al., Development and validation of the Wisconsin Smoking Withdrawal Scale. Exp Clin Psychopharmacol, 1999. 7(4): p. 354–61.

52. Zhang, L., et al., Withdrawal from chronic nicotine exposure alters dopamine signaling dynamics in the nucleus accumbens. Biological psychiatry, 2012. 71(3): p. 184–191.

53. Robinson, J.D., et al., A multimodal approach to assessing the impact of nicotine dependence, nicotine abstinence, and craving on negative affect in smokers. Exp Clin Psychopharmacol, 2011. 19(1): p. 40–52.

54. Ríos-Bedoya, C.F., et al., Association of withdrawal features with nicotine dependence as measured by the Fagerström Test for Nicotine Dependence (FTND). Addictive behaviors, 2008. 33(8): p. 1086–1089.

55. Baker, T.B., et al., Are tobacco dependence and withdrawal related amongst heavy smokers? Relevance to conceptualizations of dependence. Journal of Abnormal Psychology, 2012. 121(4): p. 909.

56. Mars, R.B., et al., Comparing brains by matching connectivity profiles. Neurosci Biobehav Rev, 2016. 60: p. 90–7.

57. Zang, Y., et al., Regional homogeneity approach to fMRI data analysis. Neuroimage, 2004. 22(1): p. 394–400.

58. Farahani, F.V., W. Karwowski, and N.R. Lighthall, Application of graph theory for identifying connectivity patterns in human brain networks: a systematic review. frontiers in Neuroscience, 2019. 13: p. 585.

59. Bassett, D.S. and O. Sporns, Network neuroscience. Nature neuroscience, 2017. 20(3): p. 353–364.

60. Johnen, V.M., et al., Causal manipulation of functional connectivity in a specific neural pathway during behaviour and at rest. Elife, 2015. 4: p. e04585.

61. Frégnac, Y., et al., Temporal covariance of pre-and postsynaptic activity regulates functional connectivity in the visual cortex. Journal of neurophysiology, 1994. 71(4): p. 1403–1421.

62. Passingham, R.E., K.E. Stephan, and R. Kötter, The anatomical basis of functional localization in the cortex. Nature Reviews Neuroscience, 2002. 3(8): p. 606–616.

63. Aertsen, A., et al., Dynamics of neuronal firing correlation: modulation of “effective connectivity”. Journal of neurophysiology, 1989. 61(5): p. 900–917.

64. Friston, K.J., Functional and effective connectivity: a review. Brain connectivity, 2011. 1(1): p. 13–36.

65. Moran-Santa Maria, M.M., et al., Right anterior insula connectivity is important for cue-induced craving in nicotine-dependent smokers. Addiction biology, 2015. 20(2): p. 407–414.

66. Wang, K., et al., The neural mechanisms underlying the acute effect of cigarette smoking on chronic smokers. PLoS One, 2014. 9(7): p. e102828.

67. Wilcox, C.E., et al., Functional network connectivity predicts treatment outcome during treatment of nicotine use disorder. Psychiatry Research: Neuroimaging, 2017. 265: p. 45–53.

68. Furlong, T.M. and L.H. Corbit, Drug addiction: augmented habit learning or failure of goal-directed control?, in Goal-directed decision making. 2018, Elsevier. p. 367–386.

69. Tricomi, E., B.W. Balleine, and J.P. O’Doherty, A specific role for posterior dorsolateral striatum in human habit learning. European Journal of Neuroscience, 2009. 29(11): p. 2225–2232.

70. Guida, P., et al., Striatal role in everyday-life and laboratory-developed habits. bioRxiv, 2021.

71. Redgrave, P., et al., Goal-directed and habitual control in the basal ganglia: implications for Parkinson’s disease. Nature Reviews Neuroscience, 2010. 11(11): p. 760–772.

72. Korponay, C., E.A. Stein, and T.J. Ross, Laterality Hotspots in the Striatum. Cereb Cortex, 2021.

73. Feng, C., et al., Individualized prediction of trait narcissism from whole-brain resting-state functional connectivity. Hum Brain Mapp, 2018. 39(9): p. 3701–3712.

74. Wang, Z., et al., Connectome-Based Predictive Modeling of Individual Anxiety. Cereb Cortex, 2021. 31(6): p. 3006–3020.

