## Supplemental Materials for "Dissociable misconfigurations of striatal functional connectivity profiles in smokers"

### Supplemental Methods

#### *Experimental Design*

Upon arrival for their MRI scanning sessions, participants underwent a medical assessment, including a urine test for recent use of illicit drugs (opiates, oxycodone, benzodiazepines, buprenorphine, cocaine, amphetamines/methamphetamines, tetrahydrocannabinol (THC), methadone, phencyclidine, and methlenedioxymethamphetamine (MDMA) and alcohol (Breathalyzer®, Alco-Sensor IV, Intoximeters Incorporated, St. Louis, MO, USA). Positive urine tests were exclusionary for all drugs except THC. For THC, positive urine tests were followed by the Drug Evaluation and Classification neuromotor exam to determine whether the participant was acutely intoxicated[1]. Positive neuromotor exams were exclusionary. Smokers were instructed to smoke *ad lib* prior to the first (sated) MR scanning session; the last cigarette was smoked an average of 48.6 min (SEM=7.69) before MR scanning[2]. Evidence of recent smoking and/or abstinence was assessed by participant self-report of last cigarette and measurement of expired carbon monoxide (CO) (BreathCO, Vitalograph, Lenexa, KS). An expired CO sample $< 5$ ppm was required to biochemically verify abstinence[3]. In the event of self-reported lapse or CO values $> 5$ ppm, the second (abstinence) MR scanning session was rescheduled[4].

#### *Curating a Matched Normative Sample*

To curate a matched normative sample, we used nearest neighbor matching via the R program MatchIt (https://cran.r-project.org/web/packages/MatchIt/MatchIt.pdf) to select healthy, non-smoking or other substance-using subjects (defined below) from the Human Connectome Project (HCP), matched to our empirical sample on age, gender, years of education, and head motion (measured by average framewise displacement on non-censored frames). Individuals from the HCP database were excluded if they reported a family history of schizophrenia, met DSM-IV criteria for alcohol dependence, reported a lifetime history of repeated substance use (>10 instances of cocaine, hallucinogen, opiate, sedatives, or stimulant use, >20 instances of tobacco use or >100 instances of marijuana use), had a urine sample on the day of scanning that was positive for any substance of abuse (cocaine, marijuana, opiates, amphetamine, or methamphetamine), or had a breath sample indicating >0.05 blood alcohol content on the day of scanning. Framewise displacement (FD) for each non-censored frame was computed by applying the following formula[5] to each subject's timeseries of rigid body parameters:

$$FD_i=|\Delta d_{ix}|+|\Delta d_{iy}|+|\Delta d_{iz}|+|\Delta \alpha_i|+|\Delta \beta_i|+|\Delta \gamma_i|$$

In all, this procedure resulted in a normative, healthy sample of 79 HCP subjects (the same total size of the empirical sample) age-, gender-, years of education-, and head motion-matched to the empirical sample.

*Resting-State fMRI Preprocessing*

Results included in this manuscript come from preprocessing performed using FMRIPREP version 20.2.1 [6, 7, RRID:SCR_016216], a Nipype [8, 9, RRID:SCR_002502] based tool. Text in below section was auto-generated via https://fmriprep.readthedocs.io/en/stable/citing.html).

Each T1w (T1-weighted) volume was corrected for INU (intensity non-uniformity) using N4BiasFieldCorrection v2.1.0 [10] and skull-stripped using antsBrainExtraction.sh v2.1.0 (using the OASIS template). Brain surfaces were reconstructed using recon-all from FreeSurfer v6.0.1 [11, RRID:SCR_001847], and the brain mask estimated previously was refined with a custom variation of the method to reconcile ANTs-derived and FreeSurfer-derived segmentations of the cortical gray-matter of Mindboggle [26, RRID:SCR_002438]. Spatial normalization to the

ICBM 152 Nonlinear Asymmetrical template version 2009c [12, RRID:SCR_008796] was performed through nonlinear registration with the antsRegistration tool of ANTs v2.1.0 [13, RRID:SCR_004757], using brain-extracted versions of both T1w volume and template. Brain tissue segmentation of cerebrospinal fluid (CSF), white-matter (WM) and gray-matter (GM) was performed on the brain-extracted T1w using fast [22] (FSL v5.0.9, RRID:SCR_002823).

Functional data was slice time corrected using 3dTshift from AFNI v16.2.07 [16, RRID:SCR_005927] and motion corrected using mcflirt (FSL v5.0.9 [14]). This was followed by co-registration to the corresponding T1w using boundary-based registration [21] with six degrees of freedom, using bbregister (FreeSurfer v6.0.1). Motion correcting transformations, BOLD-to-T1w transformation and T1w-to-template (MNI) warp were concatenated and applied in a single step using antsApplyTransforms (ANTs v2.1.0) using Lanczos interpolation.

Physiological noise regressors were extracted applying CompCor [23]. Principal components were estimated for the two CompCor variants: temporal (tCompCor) and anatomical (aCompCor). A mask to exclude signal with cortical origin was obtained by eroding the brain mask, ensuring it only contained subcortical structures. Six tCompCor components were then calculated including only the top 5% variable voxels within that subcortical mask. For aCompCor, six components were calculated within the intersection of the subcortical mask and the union of CSF and WM masks calculated in T1w space, after their projection to the native space of each functional run. Frame-wise displacement [24] was calculated for each functional run using the implementation of Nipype.

Many internal operations of FMRIPREP use Nilearn [27, RRID:SCR_001362], principally within the BOLD-processing workflow. For more details of the pipeline see https://fmriprep.readthedocs.io/en/20.2.1/workflows.html.

*Regions of Interest*

Regions of interest (ROIs) used for generation of voxel-wise striatal fingerprints were based on the frontal cortical parcellation definitions from the Harvard-Oxford cortical atlas (**Supplementary Figure 1**).

Figure S1.

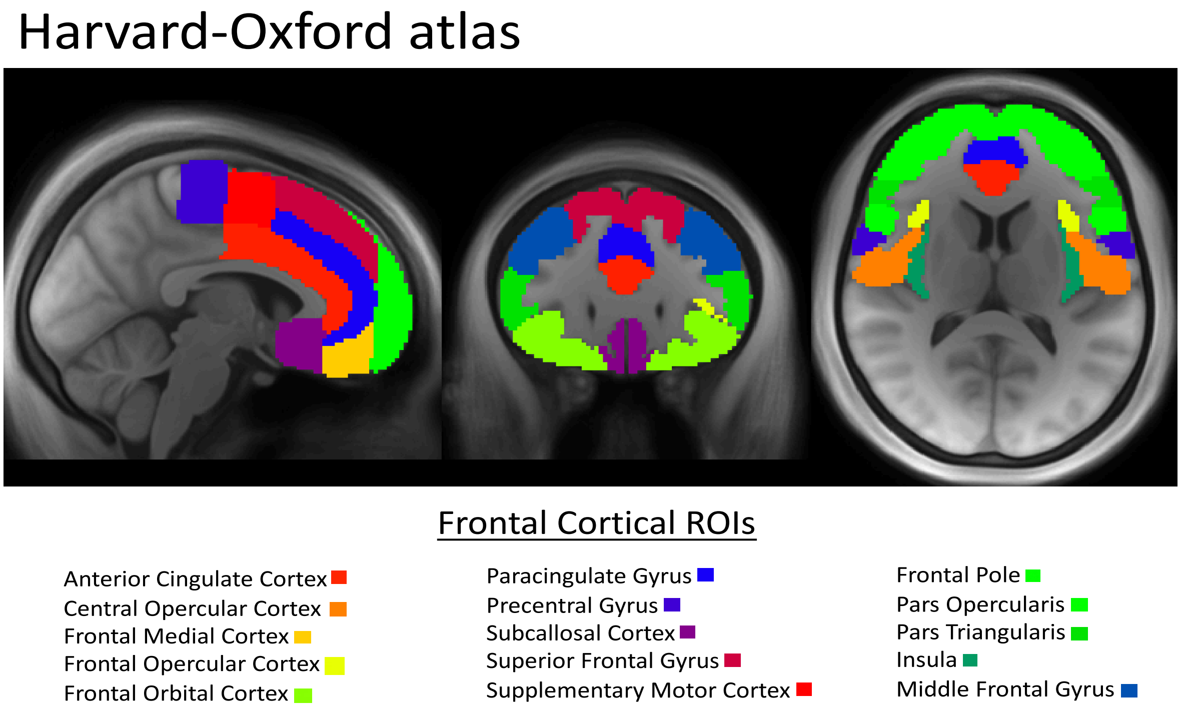


Figure S1. Frontal cortical regions of interest (ROIs) in the Harvard-Oxford atlas used as “targets” in the striatal connectivity profiles.

*Head Motion*

To examine the potential for residual motion to affect FC Z-score maps even after motion censoring, we assessed group/condition differences in subject average FD on non-censored frames. Average subject FD was higher in non-smokers than in smokers in the sated state, $t(77)=2.13, p=0.036$, and higher in smokers in the abstinent state than in the sated state, $t(45)=5.478, p<0.001$; average subject FD was not significantly different between non-smokers and smokers in the abstinent state, $t(77)=-1.74, p=0.085$ (**Supplementary Figure 2a**). The number of frames censored differed significantly between smokers in the sated (mean = 7.54) and abstinent (mean = 13.96) states, t(45) = 3.082, p=0.0035, but not between non-smokers (mean = 12.24) and smokers in the sated state, t(77)=1.907, p=0.060, or smokers in the abstinent state, t(77)=0.542, p=0.589 (**Supplementary Figure 2b**).

Figure S2.

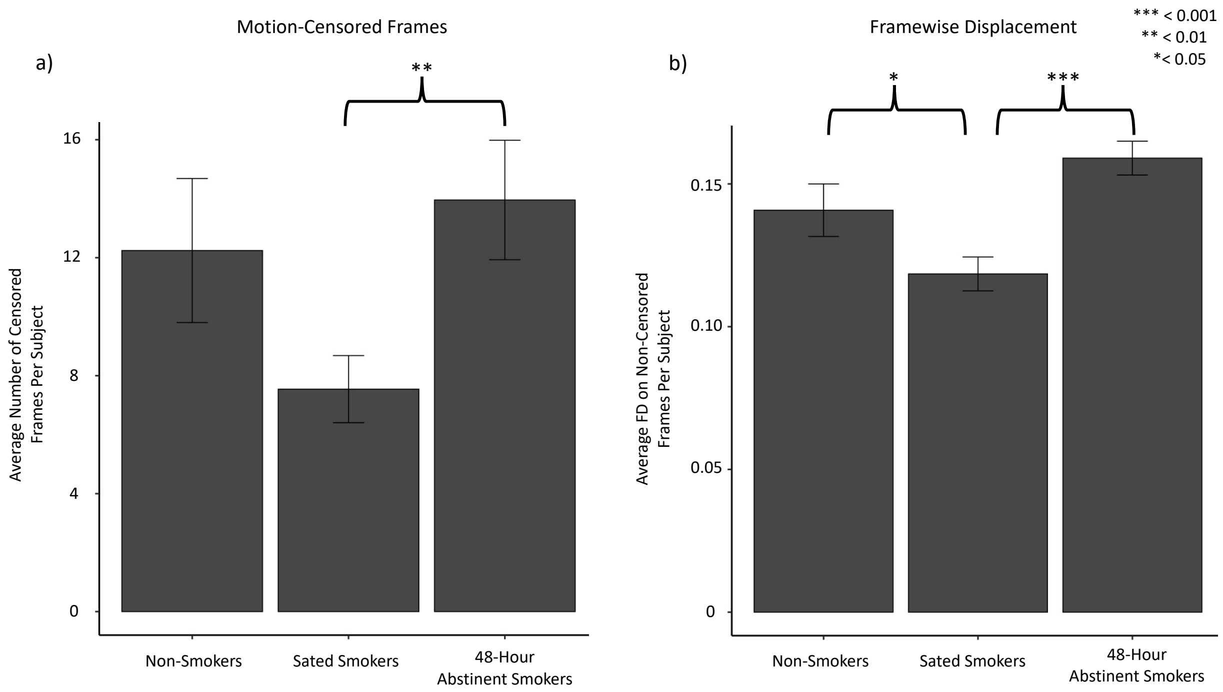


Figure S2. a) Group means for average number of motion-censored frames per subject. B) Groups means for average framewise displacement on non-motion-censored frames per subject.

Given these group differences in motion, we FD-adjusted each subject's FC Z-score maps. To do this, we first regressed average subject FD on non-censored frames against the FC Z-score map for each ROI across all subjects and computed residual maps for each ROI for each subject, in order to isolate the components of FC not attributable to variance in FD. We then added the group average FC Z-score map for each ROI back onto each subject's respective ROI residual maps to regain scale. These FD-adjusted Z-score maps served as the inputs to all main analyses.

Main analyses entailed the computation of three separate connectivity reconfiguration metrics, whose calculation is schematized in **Supplementary Figure 3**.

Figure S3.

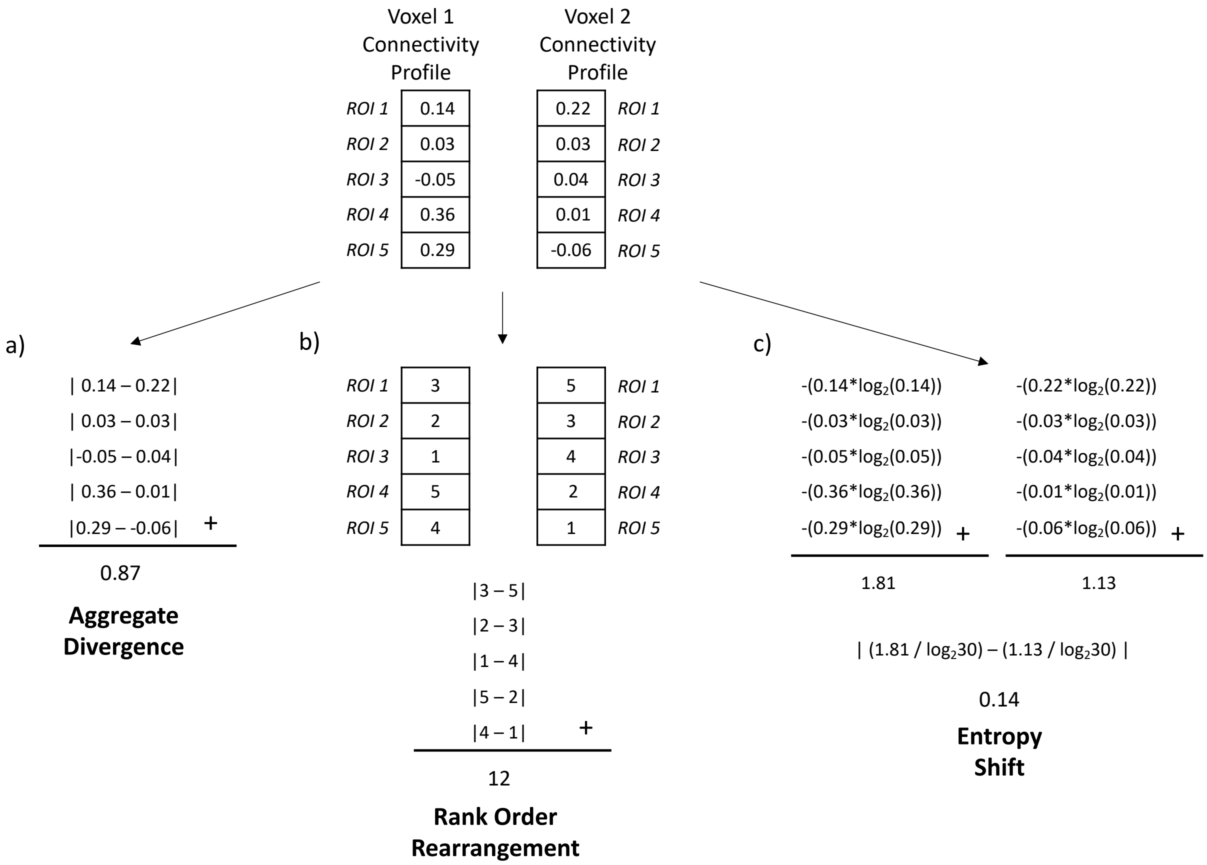


Figure S3. Illustration of computation procedure for calculating the a) aggregate divergence, b) rank order rearrangement, and c) entropy shift between the connectivity profiles of two voxels with five target ROIs. Numbers in the connectivity profile represent Z-scored functional connectivity between the voxel and each target ROI.

**Supplementary Figure 4** more concretely illustrates the voxel-wise computation of aggregate divergence between smoker and non-smoker Z-scored striatal functional connectivity maps.

Figure S4.

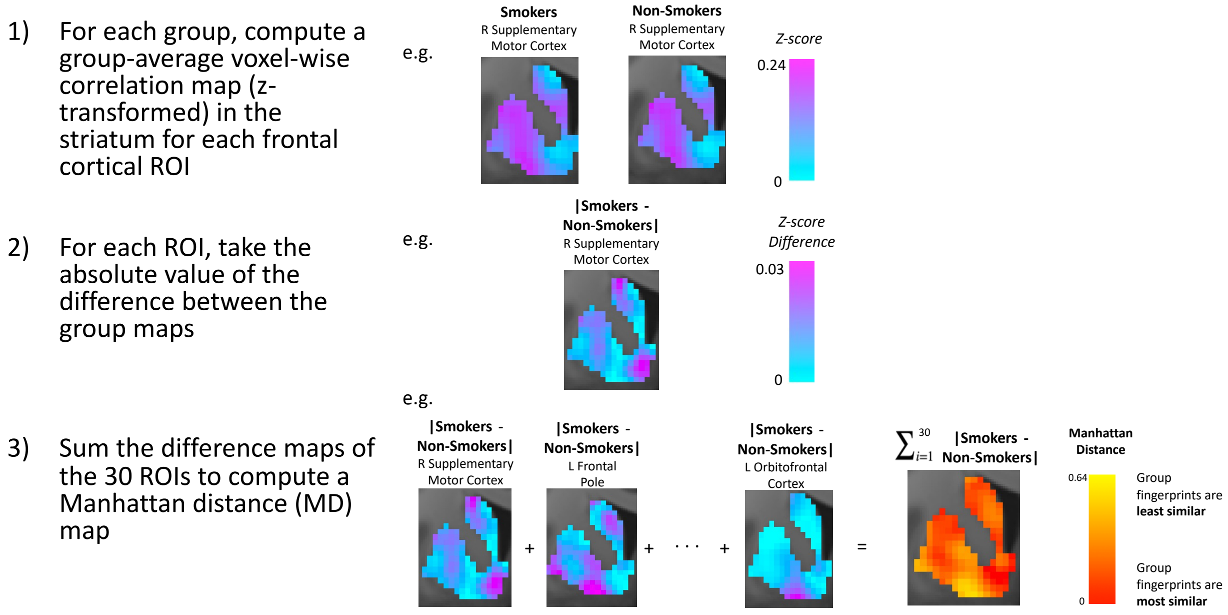


Figure S4. Illustration of computation procedure for calculating voxel-wise aggregate divergence between smoker and non-smoker groups from Z-scored functional connectivity maps.

**Supplementary Figure 5** displays the normative distributions and voxel-wise p<0.001 significance thresholds of the three connectivity profile reconfiguration metrics computed from the permutation procedure using the matched HCP sample.

Figure S5.

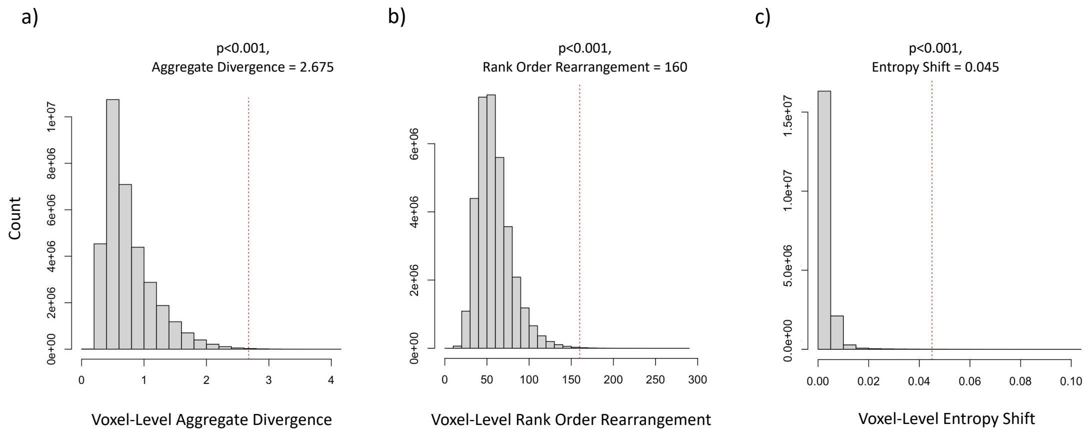


Figure S5. Normative distributions and voxel-wise p<0.001 thresholds for the connectivity profile reconfiguration metrics.

### Supplemental Results

Figure S6.

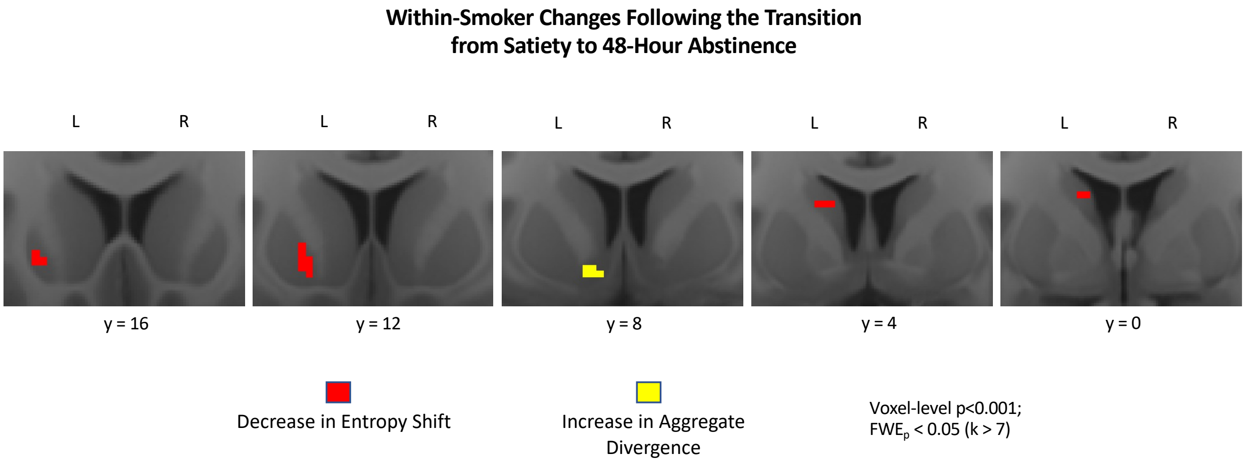


Figure S6. Clusters of significant connectivity profile reconfiguration within smokers following the transition from nicotine-satiety to 48 hour abstinence.

Figure S7.

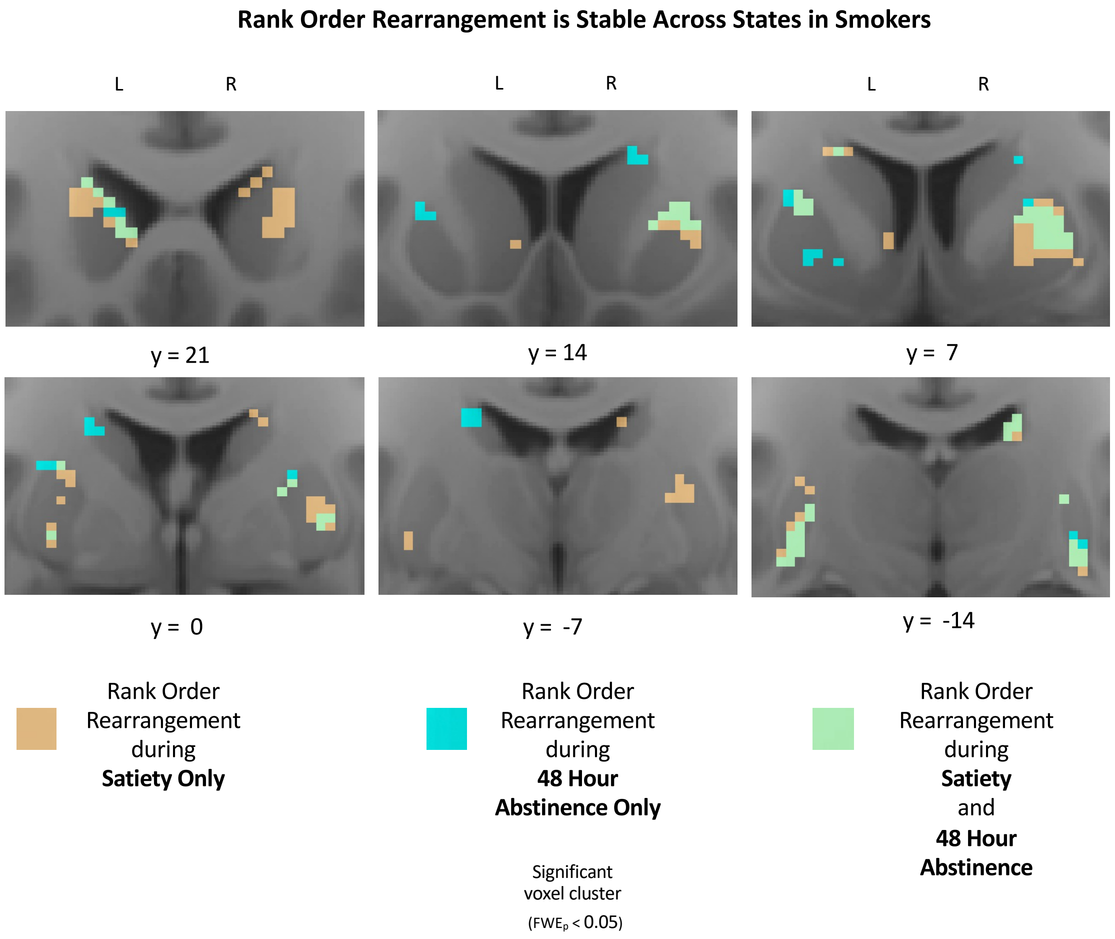


Figure S7. Clusters of significant rank order rearrangement in sated smokers (brown) and acutely abstinent smokers (blue) compared to non-smokers. Green clusters indicate areas where significant rank order rearrangement was present during both satiety and acute abstinence.

Figure S8.

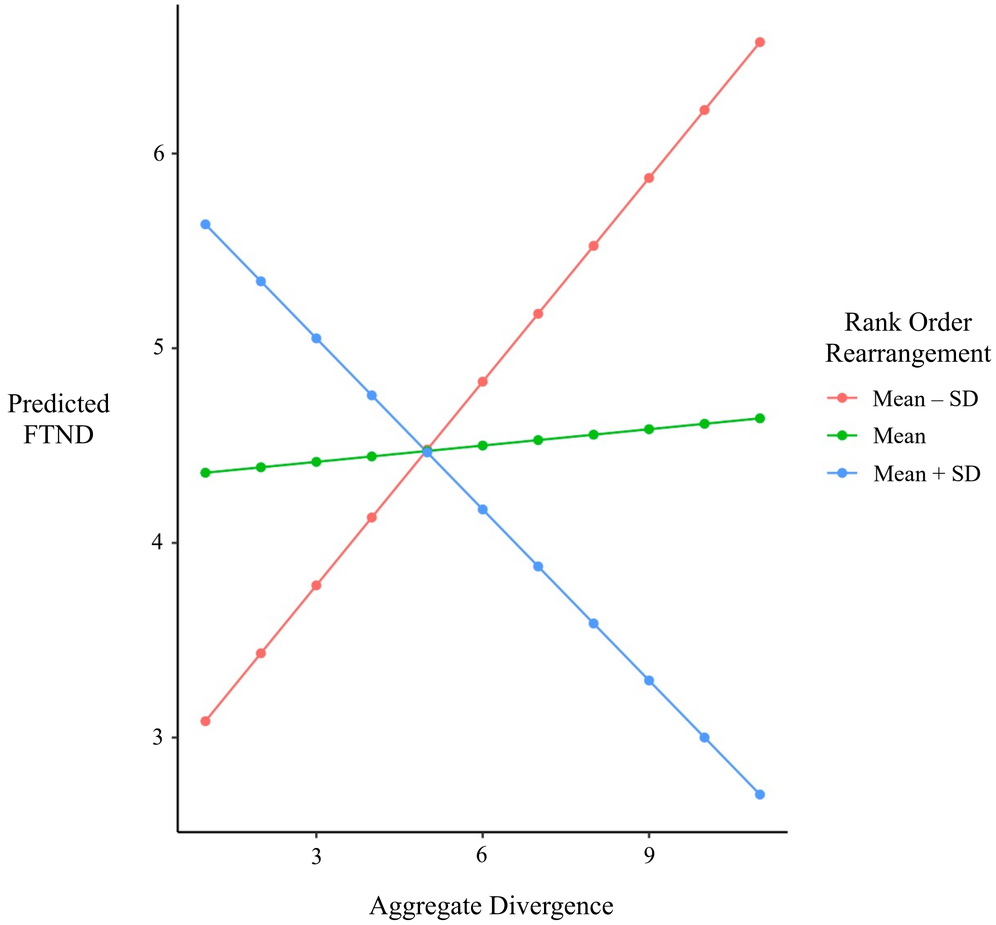


Figure S8. Unpacking the interaction between aggregate divergence and rank order rearrangement in the right caudal ventral putamen. The relationship between aggregate divergence and dependence severity (indexed by FTND score) is shown for three representative levels of rank order rearrangement: the mean minus one standard deviation (red), the mean (green), and the mean plus one standard deviation (blue). The model predicts low FTND when both metrics are high (lower right quadrant) and when both metrics are low (lower left quadrant). The model predicts high FTND when one metric is high and when one metric is low (top quadrants).

Figure S9.

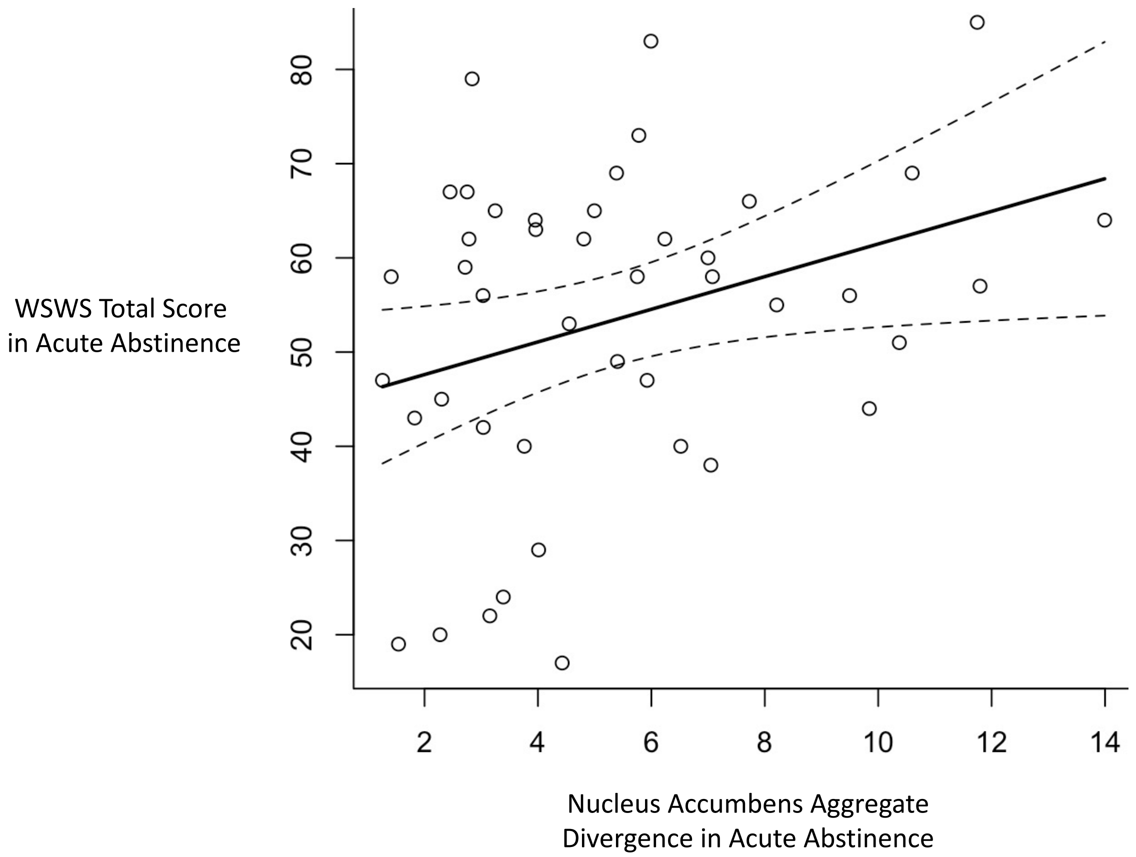


Figure S9. Relationship between aggregate divergence in the left nucleus accumbens of acutely abstinent smokers and withdrawal symptomology (indexed by WSWS total score) during acute abstinence.

**References**

1. Heishman, S.J., E.G. Singleton, and D.J. Crouch, Laboratory validation study of drug evaluation and classification program: ethanol, cocaine, and marijuana. J Anal Toxicol, 1996. **20**(6): p. 468-83.

2. Fedota, J.R., et al., Insula demonstrates a non-linear response to varying demand for cognitive control and weaker resting connectivity with the executive control network in smokers. Neuropsychopharmacology, 2016. **41**(10): p. 2557-2565.

3. Javors, M.A., J.P. Hatch, and R.J. Lamb, *Cut-off levels for breath carbon monoxide as a marker for cigarette smoking.* Addiction, 2005. **100**(2): p. 159-167.

4. Fedota, J.R., et al., *Nicotine abstinence influences the calculation of salience in discrete insular circuits.* Biological Psychiatry: Cognitive Neuroscience and Neuroimaging, 2018. **3**(2): p. 150-159.

5. Power, J.D., et al., Spurious but systematic correlations in functional connectivity MRI networks arise from subject motion. Neuroimage, 2012. 59(3): p. 2142-54.

6. Esteban O, Markiewicz CJ, Blair RW, Moodie CA, Isik AI, Erramuzpe A, Kent JD, Goncalves M, DuPre E, Snyder M, Oya H, Ghosh SS, Wright J, Durnez J, Poldrack RA, Gorgolewski KJ. fMRIPrep: a robust preprocessing pipeline for functional MRI. Nat Meth. 2018; doi:10.1038/s41592-018-0235-4

7. fMRIPrep Available from: 10.5281/zenodo.852659.

8. Gorgolewski K, Burns CD, Madison C, Clark D, Halchenko YO, Waskom ML, Ghosh SS. Nipype: a flexible, lightweight and extensible neuroimaging data processing framework in python. Front Neuroinform. 2011 Aug 22;5(August):13. doi:10.3389/fninf.2011.00013.

9. Gorgolewski KJ, Esteban O, Ellis DG, Notter MP, Ziegler E, Johnson H, Hamalainen C, Yvernault B, Burns C, Manhães-Savio A, Jarecka D, Markiewicz CJ, Salo T, Clark D, Waskom M, Wong J, Modat M, Dewey BE, Clark MG, Dayan M, Loney F, Madison C, Gramfort A, Keshavan A, Berleant S, Pinsard B, Goncalves M, Clark D, Cipollini B, Varoquaux G, Wassermann D, Rokem A, Halchenko YO, Forbes J, Moloney B, Malone IB, Hanke M, Mordom D, Buchanan C, Pauli WM, Huntenburg JM, Horea C, Schwartz Y, Tungaraza R, Iqbal S, Kleesiek J, Sikka S, Frohlich C, Kent J, Perez-Guevara M, Watanabe A, Welch D, Cumba C, Ginsburg D, Eshaghi A, Kastman E, Bougacha S, Blair R, Acland B, Gillman A, Schaefer A, Nichols BN, Giavasis S, Erickson D, Correa C, Ghayoor A, Küttner R, Haselgrove C, Zhou D, Craddock RC, Haehn D, Lampe L, Millman J, Lai J, Renfro M, Liu S, Stadler J, Glatard T, Kahn AE, Kong X-Z, Triplett W, Park A, McDermottroe C, Hallquist M, Poldrack R, Perkins LN, Noel M, Gerhard S, Salvatore J, Mertz F, Broderick W, Inati S, Hinds O, Brett M, Durnez J, Tambini A, Rothmei S, Andberg SK, Cooper G, Marina A, Mattfeld A, Urchs S, Sharp P, Matsubara K, Geisler D, Cheung B, Floren A, Nickson T, Pannetier N, Weinstein A, Dubois M, Arias J, Tarbert C, Schlamp K, Jordan K, Liem F, Saase V, Harms R, Khanuja R, Podranski K, Flandin G, Papadopoulos Orfanos D, Schwabacher I, McNamee D, Falkiewicz M, Pellman J,

Linkersdörfer J, Varada J, Pérez-García F, Davison A, Shachnev D, Ghosh S. Nipype: a flexible, lightweight and extensible neuroimaging data processing framework in Python. 2017. doi:10.5281/zenodo.581704.

10. Tustison NJ, Avants BB, Cook PA, Zheng Y, Egan A, Yushkevich PA, Gee JC. N4ITK: improved N3 bias correction. IEEE Trans Med Imaging. 2010 Jun;29(6):1310–20. doi:10.1109/TMI.2010.2046908.

11. Dale A, Fischl B, Sereno MI. Cortical Surface-Based Analysis: I. Segmentation and Surface Reconstruction. Neuroimage. 1999;9(2):179–94. doi:10.1006/nimg.1998.0395.

12. Fonov VS, Evans AC, McKinstry RC, Almli CR, Collins DL. Unbiased nonlinear average age-appropriate brain templates from birth to adulthood. NeuroImage; Amsterdam. 2009 Jul 1;47:S102. doi:10.1016/S1053-8119(09)70884-5.

13. Avants BB, Epstein CL, Grossman M, Gee JC. Symmetric diffeomorphic image registration with cross-correlation: evaluating automated labeling of elderly and neurodegenerative brain. Med Image Anal. 2008 Feb;12(1):26–41. doi:10.1016/j.media.2007.06.004.

14. Jenkinson M, Bannister P, Brady M, Smith S. Improved optimization for the robust and accurate linear registration and motion correction of brain images. Neuroimage. 2002 Oct;17(2):825–41. doi:10.1006/nimg.2002.1132.

15. Andersson JLR, Skare S, Ashburner J. How to correct susceptibility distortions in spin-echo echo-planar images: application to diffusion tensor imaging. Neuroimage. 2003 Oct;20(2):870–88. doi:10.1016/S1053-8119(03)00336-7.

16. Cox RW. AFNI: software for analysis and visualization of functional magnetic resonance neuroimages. Comput Biomed Res. 1996 Jun;29(3):162–73. doi:10.1006/cbmr.1996.0014.

17. Jenkinson M. Fast, automated, N-dimensional phase-unwrapping algorithm. Magn Reson Med. 2003 Jan;49(1):193–7. doi:10.1002/mrm.10354.

18. Huntenburg JM. Evaluating nonlinear coregistration of BOLD EPI and T1w images. Freie Universität Berlin; 2014. Available from: http://hdl.handle.net/11858/00-001M-0000-002B-1CB5-A.

19. Wang S, Peterson DJ, Gatenby JC, Li W, Grabowski TJ, Madhyastha TM. Evaluation of Field Map and Nonlinear Registration Methods for Correction of Susceptibility Artifacts in Diffusion MRI. Front Neuroinform. 2017 [cited 2017 Feb 21];11. doi:10.3389/fninf.2017.00017.

20. Treiber JM, White NS, Steed TC, Bartsch H, Holland D, Farid N, McDonald CR, Carter BS, Dale AM, Chen CC. Characterization and Correction of Geometric Distortions in 814 Diffusion Weighted Images. PLoS One. 2016 Mar 30;11(3):e0152472. doi:10.1371/journal.pone.0152472.

21. Greve DN, Fischl B. Accurate and robust brain image alignment using boundary-based registration. Neuroimage. 2009 Oct;48(1):63–72. doi:10.1016/j.neuroimage.2009.06.060.

22. Zhang Y, Brady M, Smith S. Segmentation of brain MR images through a hidden Markov random field model and the expectation-maximization algorithm. IEEE Trans Med Imaging. 2001 Jan;20(1):45–57. doi:10.1109/42.906424.

23. Behzadi Y, Restom K, Liau J, Liu TT. A component based noise correction method (CompCor) for BOLD and perfusion based fMRI. Neuroimage. 2007 Aug 1;37(1):90–101. doi:10.1016/j.neuroimage.2007.04.042.

24. Power JD, Mitra A, Laumann TO, Snyder AZ, Schlaggar BL, Petersen SE. Methods to detect, characterize, and remove motion artifact in resting state fMRI. Neuroimage. 2013 Aug 29;84:320–41. doi:10.1016/j.neuroimage.2013.08.048.

25. Pruim RHR, Mennes M, van Rooij D, Llera A, Buitelaar JK, Beckmann CF. ICA-AROMA: A robust ICA-based strategy for removing motion artifacts from fMRI data. Neuroimage. 2015 May 15;112:267–77. doi:10.1016/j.neuroimage.2015.02.064.

26. Klein A, Ghosh SS, Bao FS, Giard J, Häme Y, Stavsky E, et al. Mindboggling morphometry of human brains. PLoS Comput Biol 13(2): e1005350. 2017. doi:10.1371/journal.pcbi.1005350.

27. Abraham A, Pedregosa F, Eickenberg M, Gervais P, Mueller A, Kossaifi J, Gramfort A, Thirion B, Varoquaux G. Machine learning for neuroimaging with scikit-learn. Front in Neuroinf 8:14. 2014. doi:10.3389/fninf.2014.00014.
